# A simulation and comparison of dynamic functional connectivity methods

**DOI:** 10.1101/212241

**Authors:** William Hedley Thompson, Craig Geoffrey Richter, Pontus Plavén-Sigray, Peter Fransson

## Abstract

There is a current interest in quantifying brain dynamic functional connectivity (DFC) based on neuroimaging data such as fMRI. Many methods have been proposed, and are being applied, revealing new insight into the brain’s dynamics. However, given that the ground truth for DFC in the brain is unknown, many concerns remain regarding the accuracy of proposed estimates. Since there exists many DFC methods it is difficult to assess differences in dynamic brain connectivity between studies. Here, we evaluate five different methods that together represent a wide spectrum of current approaches to estimating DFC (sliding window, tapered sliding window, temporal derivative, spatial distance and jackknife correlation). In particular, we were interested in each methods’ ability to track changes in covariance over time, which is a key property in DFC analysis. We found that all tested methods correlated positively with each other, but there were large differences in the strength of the correlations between methods. To facilitate comparisons with future DFC methods, we propose that the described simulations can act as benchmark tests for evaluation of methods. In this paper, we present *dfcbenchmarker*, which is a Python package where researchers can easily submit and compare their own DFC methods to evaluate its performance.

## Introduction

Dynamic functional connectivity (DFC) is being applied to an increasing number of topics studying the brain’s networks. Topics that have been explored with DFC include development (1), various pathologies (2,3), affect (4), attention (5), levels of consciousness (6), and temporal properties of the brain’s networks (7–9). There are many concerns raised regarding methodological issues. These issues include from biased variance (10,11), movement artefacts (12), and statistics (13,14).

Methods used to derive DFC estimates are as diverse as its range of applications. Examples of different methods include: the sliding window method, sometimes tapered (15), temporal derivatives (16), methods using Euclidean distance between spatial configurations (8), k-means clustering methods (7,17), eigenconnectivities (18), point process methods (19,20), Kalman filters (21,22). flexible least squares (23), temporal ICA (24), sliding window ICA (25), dynamic conditional correlation (26), phase differences (27) wavelet coherence (4), hidden Markov models (28), and variational Bayes hidden Markov models (29). This list of DFC methods is not exhaustive, and even more methods can be found in the literature.

While these methods and their applications may offer new insights into the functions of the brain and cognition, it becomes difficult to compare results when different studies use different methods to estimate brain dynamics. Each method is often introduced and evaluated by the authors’ own simulations, empirical demonstrations, and/or theoretical argumentations. However, apparent differences in dynamic connectivity in different studies may have been influenced, or even caused, by differences in the underlying methodology used to derive connectivity estimates.

In order to maximize reproducibility of reported findings, it is important that comparisons of proposed DFC methods can be made with a common set of simulations. To this end, we have developed four simulations that aim to show how well results from different DFC methods correlate with each other and evaluate their performance of tracking time varying covariance. The proposed methods and simulations are packaged in the Python package *dfcbenchmarker*, (available at github.com/wiheto/dfcbenchmarker). Researchers can evaluate their own DFC methods in *dfcbenchmarker*. The software also allows for new methods to be submitted to us for inclusion in future reports. Here we demonstrate the functionality and results obtained by *dfcbenchmarker* by evaluating the performance of the following five methods: sliding window (SW), tapered sliding window (TSW), spatial distance (SD), jackknife correlation (JC), and temporal derivatives (TD).

## Methods section

### Software used

All methods for DFC derivation were implemented in Teneto v0.2.1 (8). Bayesian statistics for evaluating performance of DFC methods were calculated in PyMC3 V3.1 (30), simulations and analysis were done using Numpy V1.13.1 (31), Scipy V0.19.1 (32), and Pandas V0.19.2. Matplotlib V2.0.2 (33) and Seaborn V0.7.1 (34) were used for figure creation.

### Dynamic functional connectivity methods

As discussed in the introduction, the list of published DFC methods that are designed to be applied to fMRI imaging data is long. In an ideal world all methods will be contrasted under the same conditions so that an evaluation of which methods that give appropriate results can be performed. However, it was not our intention to provide a complete comparison of all published methods. Instead we have made all simulation tools freely available so that researchers can evaluate their own DFC methods. Before describing the simulations and the results, we provide a brief overview of the five methods that are evaluated in this article.

### Sliding window (SW)

The SW method is one of the most commonly used methods to estimate DFC. The sliding window method uses a continuous subsection of the data, estimates the degree of correlation (Pearson correlation), slides the window one step in the time series, and repeats. This creates a smooth connectivity time series as neighbouring estimates of connectivity share all but two data points. The SW method is based on the assumption that nearby temporal points are helpful to estimate the covariance. In our simulations, the window length was set to 63, which is reasonable for a sliding window analysis. Given the common choice of a time resolution (TR) of 2 seconds in fMRI, this results in a window length of 126 seconds which is reasonable, given rule-of-thumb choices for window length estimates that has recently been proposed (35).

### Tapered sliding window (TSW)

The TSW method can be described as a weighted Pearson correlation where the weights are set to zero except for the data points residing inside the window. This procedure is identical to the SW method except that a larger weight is placed on time points closer to the centre of the window (*t*). Often, the weights are distributed according to a Gaussian distribution centred at *t*. In our simulations using the TSW method, we used a Gaussian distribution with a variance of 10.

The window length was the same as for the SW method (centred at *t*). See also (15) for an example of usage of the TSW method.

### Spatial Distance (SD)

An alternative to using temporally adjacent data points is to estimate the covariance using data points that have similar spatial profiles. In this case, the weights are based on distances in spatial dimensions of data acquired at different time points. In functional neuroimaging data each spatial dimension corresponds to a region of interest or voxels. One way to accomplish this is by first computing weights for each time point (*t*) to all other time points. This can be done by taking the inverse of their Euclidean distance to each other time point (*u*):

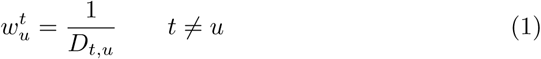

where *D* is the Euclidean distance. Hence, the closer in space the data points are located, the larger their individual weights will be. Each time point gets a weight vector (*w*_*t*_) which is the length of the time series. The weight vector for each time point are subsequently scaled between 0 and 1. The “self weights” 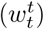 are set to 1. The connectivity estimate at *t* is the weighted Pearson correlation where each time point is weighted by *w^t^*. For more details of the SD method, see (8).

### Jackknife Correlation (JC)

The JC method has previously been shown on electrocorticographic data for signal trial coherence and Granger causality (36). To the best of our knowledge, the jackknife correlation method has not yet been utilized in the DFC literature. Thus, we provide a more detailed description of its logic and workings. The jackknife correlation method is outlined in more detail and contrasted to a binning approach (which is akin to the sliding window method) in (36). The JC method, when applied to single time point covariance estimates of signals *x* and *y* at *t* computes the Pearson correlation between the two signals using all time points in *x* and *y* with the exception of *x*_*t*_ and *y*_*t*_:

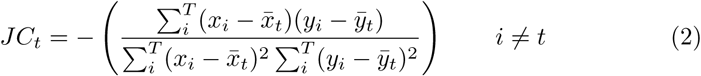

Of note, the inclusion of the minus sign in the equation above is to correct for the inversion caused by the leave-n-out process (see below). The 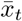 and 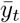 are the expected values, excluding data at time point *t*:

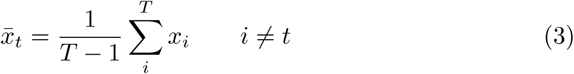

To demonstrate the JC method, 10,000 time points were drawn from a multivariate Gaussian distribution with a mean of 0 and a variance of 1 to generate the two time series shown in Figure 1A. Additionally, the time series were constructed so that the covariance between the two varied as a function of time. For the first 2,000 time points, the covariance was set to 0.8 and then further decreased in steps of 0.2 for every 2,000th time point (Figure 1B).

The *relative* connectivity time series are similar (but inverted) for a leave-*n*- out compared to a window-length-*n* methods (Figure 1C). The JC method corresponds to the case when *n*=1, i.e. a leave-1-out approach after correcting for the inversion (all leave-*n*-out estimates are multiplied by -1 to correct for the inversion in Figure 1). All possible choices of *n* were computed for both methods (i.e. a leave-1-out to a leave-9,999-out and a window-of-length-2 to window-length-10,000 (see Figures 1DE). As shown in Figure 1G, the Spearman correlation between the two methods is close to 1 for various choice of *n*. However, their correspondence in covariance estimates between the two methods departs at the tails (Figures 1F and 1H). These deviations occur for two reasons: (1) when *n* is very small it implies there is little data to work with for the window-length-*n* method. On the other hand, low values of *n* do not hamper the performance of the leave-*n*-out method. (2) Large values of *n* will result in few estimates for covariance for each time point, which makes the correlation between the two methods less stable. In sum, while it is impossible to create estimates for window-of-length-1, it is however possible to use a leave-1-out method as an approximation for a window-length-1 due to the symmetry between the two methods.

When estimating the DFC, the two major aims are to accurately measure the covariance and to be sensitive to changes in the covariance. In the case of the leave-1-out (i.e. the jackknife correlation) approach, we achieve a unique connectivity estimate per time point that is more reliable than using a smaller window size (due to the fact that more data is used). Usually, the SW method has to find a balance between the two aims. In this respect, the JC method is an optimal sliding window method as it does not have to compromise between sensitivity on one hand and accuracy on the other.

The time point based DFC estimate obtained with the JC method should be interpreted as the relative difference in connectivity at any particular data point compared to all other data points in the time series. This is because the covariance for each data point is estimated based on its relationship to all other data points. To illustrate this effect, consider the 49 data points randomly sampled from a Gaussian distribution with a mean of 0 and a covariance of 0.5 as shown in Figure 1I. If we assume that the 49 time points are used to compute the JC estimate for the covariance for a 50th time point, the value of this new data point will have no impact on its JC covariance estimate because the other 49 points are used. What the 50th time point does is change the JC estimate for the other 49 points. This means that the relative position of the 50th point changes in relation to the rest of the time series. The standardized JC estimate of covariance for all possible values of the 50th time point is shown in Figure 1J. In sum, individual JC estimates have little meaning by themselves but they do show which time points have an increased connectivity and which that have a decreased connectivity relative to all the other points in the time series.

**Figure 1:**
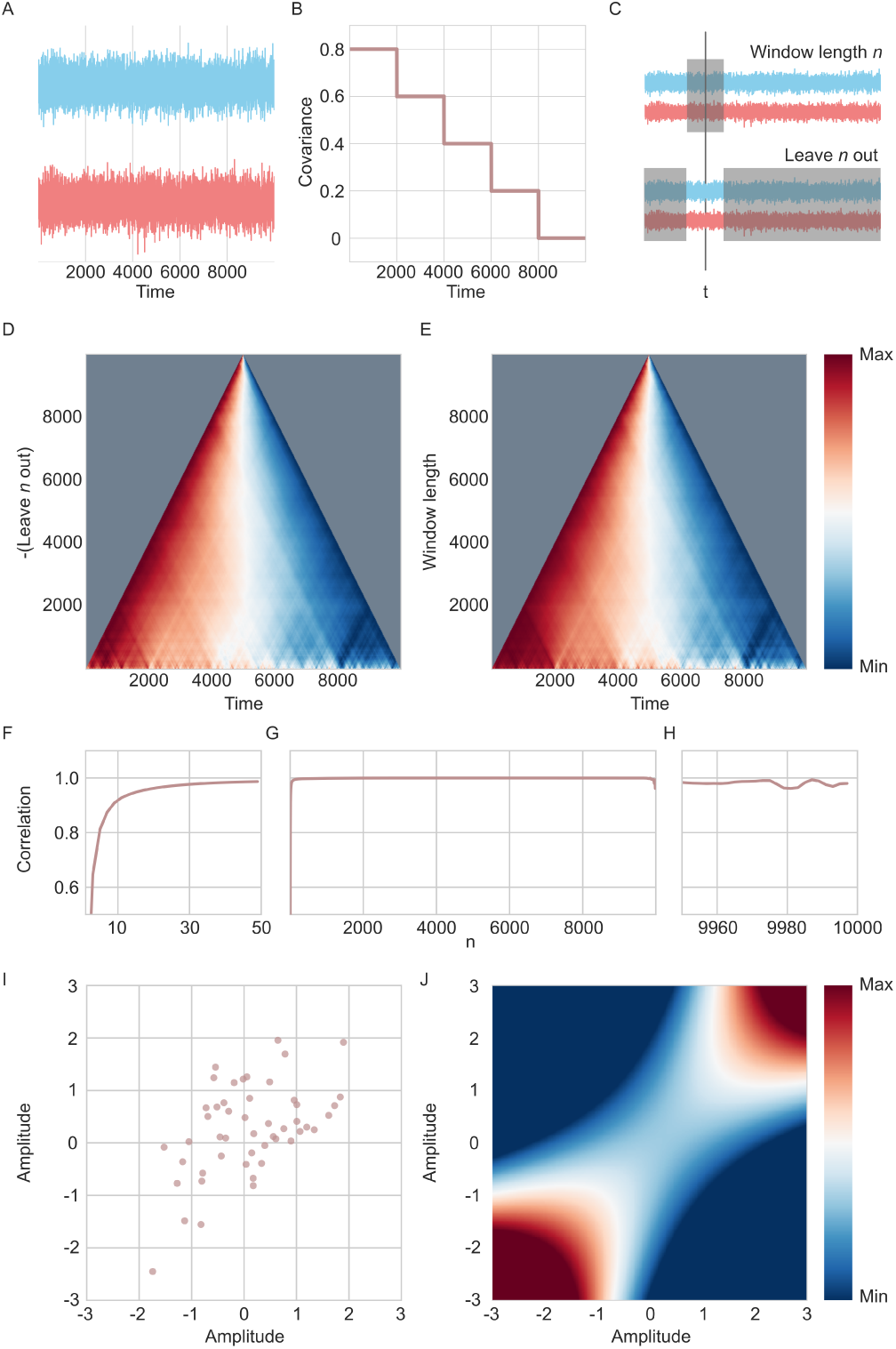
Illustration of jackknife correlation and leave-*n*-out/window-length-*n* symmetry. (A) Two time series drawn from a multi-variate Gaussian distribution, stretching over 10,000 time points with their covariance parameter changing every 2,000th time point. (B) The covariance parameter of the two time series in A. (C) Depiction of how the window-length-n and leave-*n*-out relate to each other. Shaded region indicates time points used in the correlation estimate at time point *t*. (D) The correlation estimate per time point for varying n of the leave-*n*-out method (correcting for the inversion by multiplying with -1). The time series for each *n* is scaled between 0 and 1. (E) Same as D, but for the window-length-*n* method. (F-H) Correlation between the time series of connectivity estimates for window-length-*n* and leave-*n*-out methods for different values of *n*. (F) Shows *n* between 1-50 (G) Shows *n* over the entire time series. (H) Shows *n* between 9,950 and 10,000. (I) The correlate of the amplitude of the two time series. 49 time points were sampled from a multivariate Gaussian distribution with a covariance of 0.5. (J) Illustration of the jackknife correlation estimate for different possible values, relative to the 49 time points in I.

When using the JC method to estimate DFC, it is important to keep in mind that it leads to a compression of the variance. Furthermore, the amount of compression is proportional to the length of the time series. It is often helpful to scale or standardize the connectivity time series derived by the JC method before any subsequent analysis. Finally, while a Pearson correlation was used in this study for the JC, it is possible to use other correlation methods such as the Spearman Rank instead.

### Temporal Derivative (TD)

The TD approach to estimate DFC was first introduced in (16). In brief, the temporal derivative method first computes the temporal derivative of a time series as:

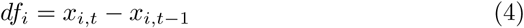

Next, the coupling between the signal sources *i* and *j* is defined as the product of the two derivatives *df*_*i*_ and *df*_*j*_ for each time point *t*, divided by the product of the standard deviation for *df*_*i*_ and *df*_*j*_:

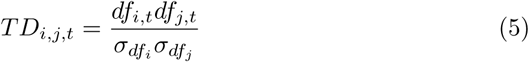

The TD method is often used together with a smoothing function in the form of a window function. In our simulations, a window length of 7 was chosen, since this was considered optimal in (16).

### Post-processing for DFC estimates

After each of the DFC methods were applied to the simulated data a Fisher transform was applied to the connectivity time series (except for the TD method). To illustrate the variance compression that results from the JC method, the DFC for the JC method was not standardized in Simulation 1.

The SW, TSW and TD methods are all effected by any autocorrelation existing in the BOLD signal, whereas the spatial distance and jackknife correlation method are not effected by any autocorrelation in the time series.

### Simulations

To compare accuracy and performance for the five DFC methods, we performed four different simulations. The first simulation investigated the similarity of the five methods by correlating their respective connectivity estimates. The second simulation targeted how well the different methods were able to track a fluctuating covariance parameter. The third simulation tested how robust the estimated fluctuating covariance is when the mean of the time series fluctuates, mimicking the haemodynamic response function. The forth simulation considered whether DFC methods can accurately track abrupt changes in covariance.

Simulations 2, 3, and 4 all consisted of fluctuating covariance parameter (*r*_*t*_) that was used to generate the time series. This is the covariance between the time series that the data was sampled from at *t*. The parameter choice for the difference simulations were considered relevant for the fMRI BOLD signal. A full account of all model assumptions made as well as a justification for our model parameter settings for the four simulations models used in the present study is given in the Supplementary Material.

### Statistics

In principle, it is possible to simply correlate the results from the different DFC methods with the *r*_*t*_ values to statistically evaluate their performance. However given the inherent, but known, uncertainty in *r*_*t*_, we deemed it was appropriate to create a statistical model which accounts for this uncertainty. Thus, for each DFC method, a Bayesian statistical model was created to evaluate the relationship between the DFC estimate and the signal covariance.

The Bayesian model aims to predict *y*, which is the vector of the known sampled covariances (i.e. *r*_*t*_) with *x*, which is the connectivity estimate for each DFC model.

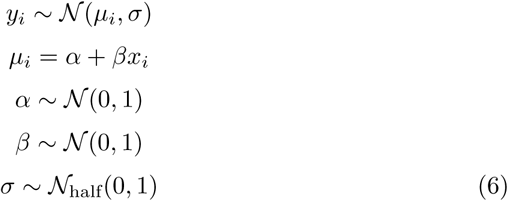

All DFC estimates and the values of *r*_*t*_ were standardized prior to calculating the models, to facilitate the interpretation of the posterior distribution parameter β. The different DFC methods vary in length of number of time points estimated (e.g. the beginning and end of the time series cannot be estimated with the sliding window method). In order to facilitate model comparison between methods, we restrained the simulations to include only the time points that had estimates from all DFC methods (i.e the limit was set by the SW and TSW methods which can estimate the covariance for 9,938 out of 10,000 time points).

The statistical model were estimated through 5,500 draws from a Markov Chain Monte Carlo (MCMC) with a No-U-Turn Sampler (37) sampler implemented in pymc3. The first 500 samples were burned.

The statistical models for the different DFC methods can be contrasted in two ways: (1) model comparison by examining the model fit; (2) by comparing the posterior distribution of β for the different DFC methods. To evaluate the model fit, the Watanabe-Akaike information criterion (WAIC) was used. The posterior distribution of β illustrates the size and uncertainty of the relationship between *x* and *y*. To aid the interpretation of these results for readers unfamiliar with Bayesian statistics, the mode of the distribution corresponds approximately to a maximum-likelihood estimated β value in a linear regression (if uniform priors are used for the parameters the posterior mode and the maximum-likelihood estimator would have been exactly the same).

## Results

### Simulation 1

The first simulation aimed to quantify the similarity of the different DFC time series estimates. If two DFC methods are strongly correlated, this is a positive sign as they are estimating similar aspects of the evolving relationship between time series. A negative correlation between two methods would suggest that they do not capture the same dynamics of the signal.

In this simulation we created two time series (*X*), each consisting of 10,000 time points in length. The time series had an autocorrelation and covariance given by:

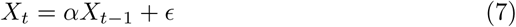

The autocorrelation with lag of 1 is determined by α*X*_*t*–1_ and the covariance at *t* is determined by *E*. *E* was sampled from a multivariate Gaussian distribution 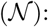

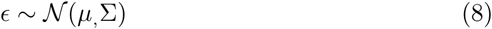

where *µ* is the mean and Σ being the covariance matrix of the multivariate Gaussian distribution. Both time series were set to have a mean of 0, variance of 1 and a covariance of 0.5. In summary:

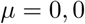

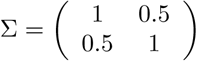

The autoregressive parameter *α* controls the size of the autocorrelation in relation to the preceding time point (i.e. the proportion of the previous time point that is kept). Here, it was set to 0.8 which was deemed to be an appropriate degree of autocorrelation for BOLD time series. A portion of the two simulated time series is found in Figure 2A together with the plots of their respective autocorrelation (Figures 2BC) and a plot of the correlation between the two time series (Figure 2D).

**Figure 2:**
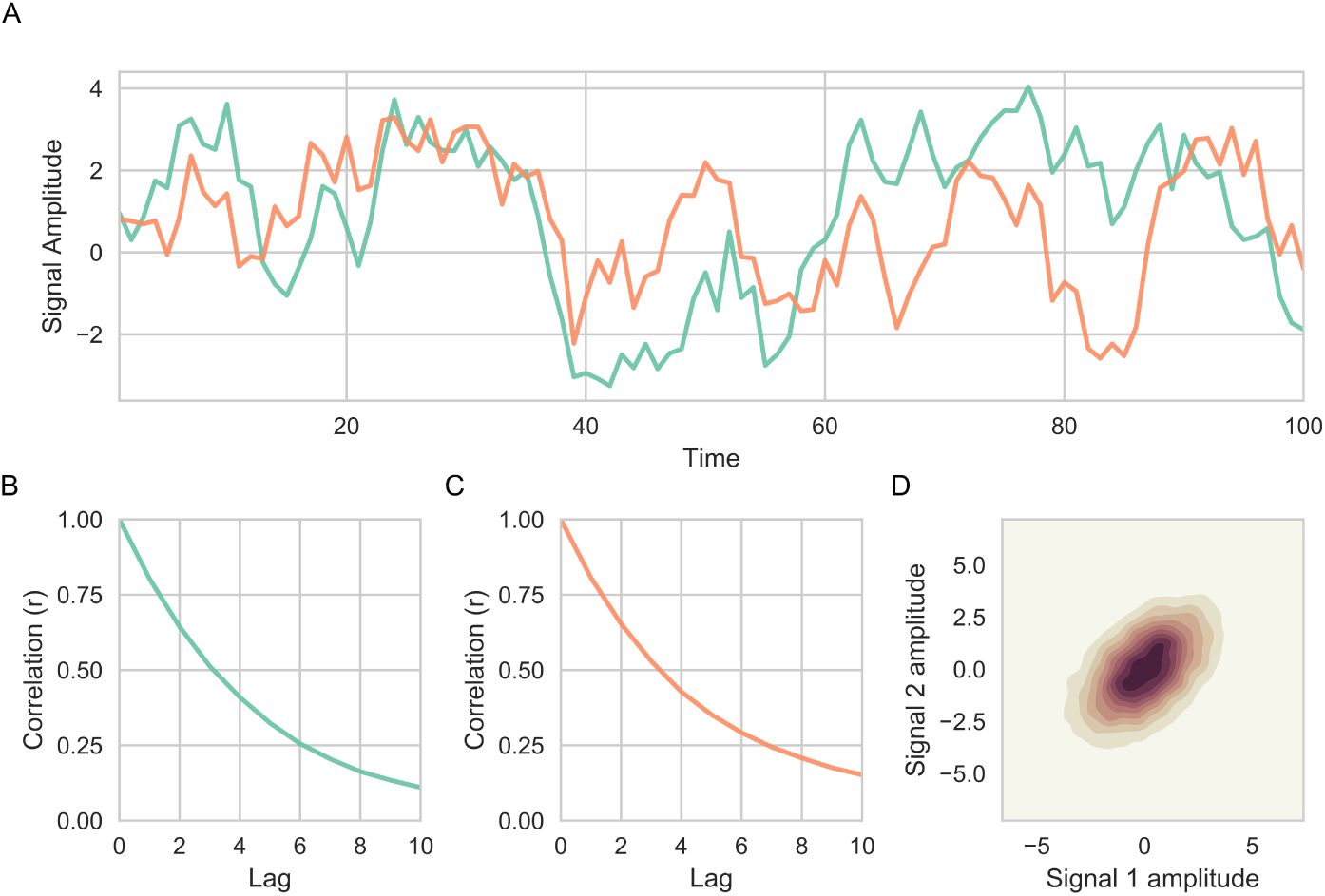
Simulated data in Simulation 1. (A) Two correlated time series were generated (a total of 10,000 time points were simulated, only the first 100 time points shown in the figure). (B-C) Spectrum of autocorrelation of both time series (colors corresponding to respective time series given in (A)) for up to 10 lags. (D) Kernel density estimation illustrating the covariance between two time series (*r* = 0.51).

The resulting connectivity time series for the five different DFC methods when applied to the simulated data is shown in Figure 3. From Figure 3, several qualitative observations can be made about the methods. Firstly, there was a very strong similarity between the SD and JC methods, despite the fact that they consist of quite different assumptions. Further, the SD, JC, and TD methods were all able to capture considerably quicker transitions than the SW and TSW methods. Finally, the variance of the JC method was considerably smaller than all other methods, illustrating the variance compression as previously discussed.

**Figure 3:**
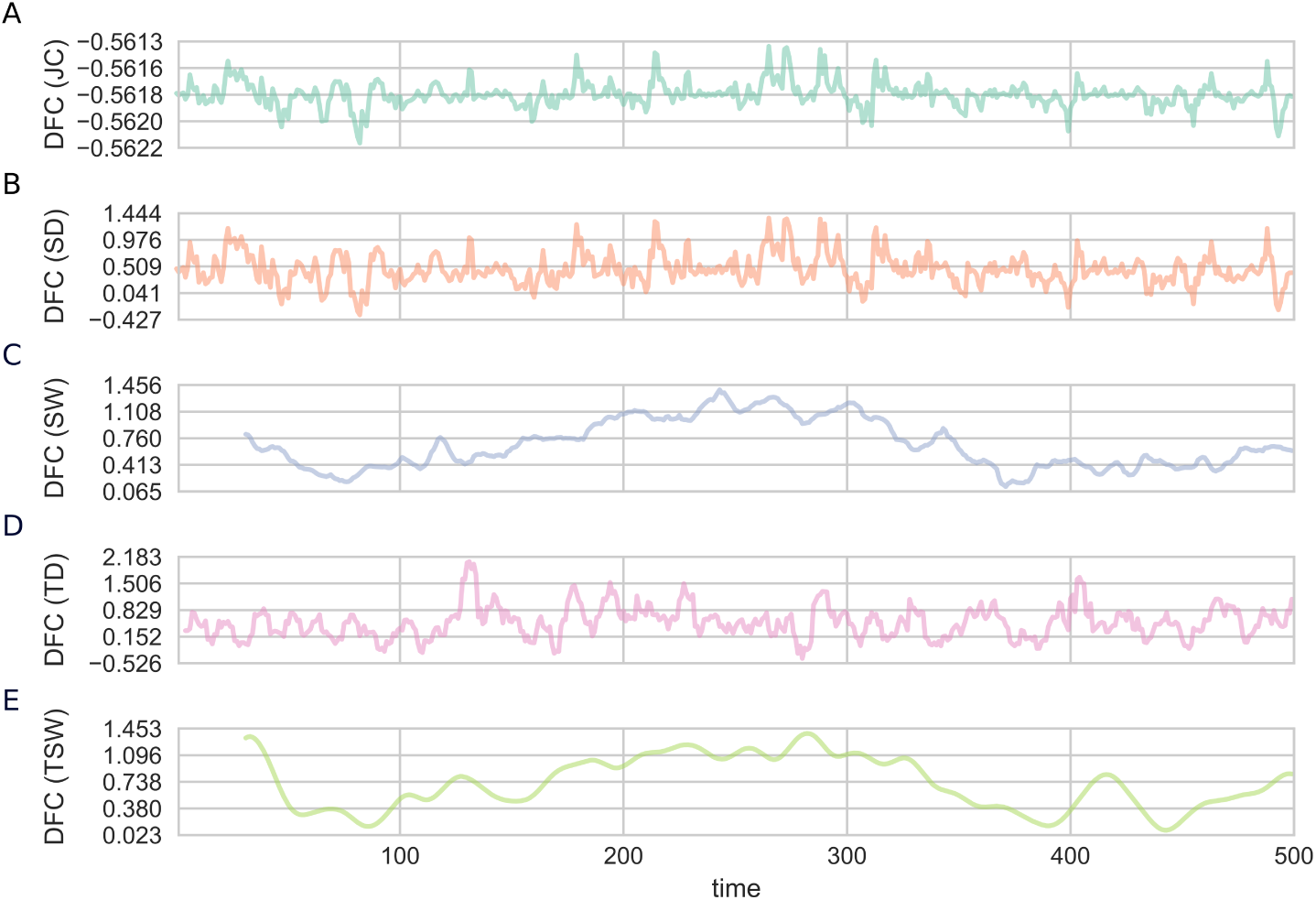
Dynamic functional connectivity estimates for Simulation 1. The five DFC methods (ordered alphabetically): (A) JC; (B) SD; (C) SW; (D) TD; (E) TSW. Only the first 500 time points are shown.

To assess the degree of similarity of the estimates of functional connectivity time series obtained from all five DFC methods, a Spearman correlations was used (Figure 4). The connectivity time series estimates from all methods correlated positively with each other (Figure 4). The SD and JC methods showed a strikingly strong relationship (ρ = 0.976). The two sliding window methods also display a strong correlation (ρ = 0.735). The lowest correlation was found between the SW and TD methods (ρ = 0.086).

The results from Simulation 1 showed that the connectivity estimates provided by the tested methods are, to a varying extent, correlated positively with each other. It also illustrated how the different methods differ in their resulting smoothness of the connectivity time series. The results from this simulation cannot validate whether any DFC method is superior to any other, it merely highlights which methods produce similar connectivity time series.

### Simulation 2

In Simulation 1, it was not possible to evaluate how well the different DFC methods perform. To evaluate the performance, the simulated data must change

**Figure 4:**
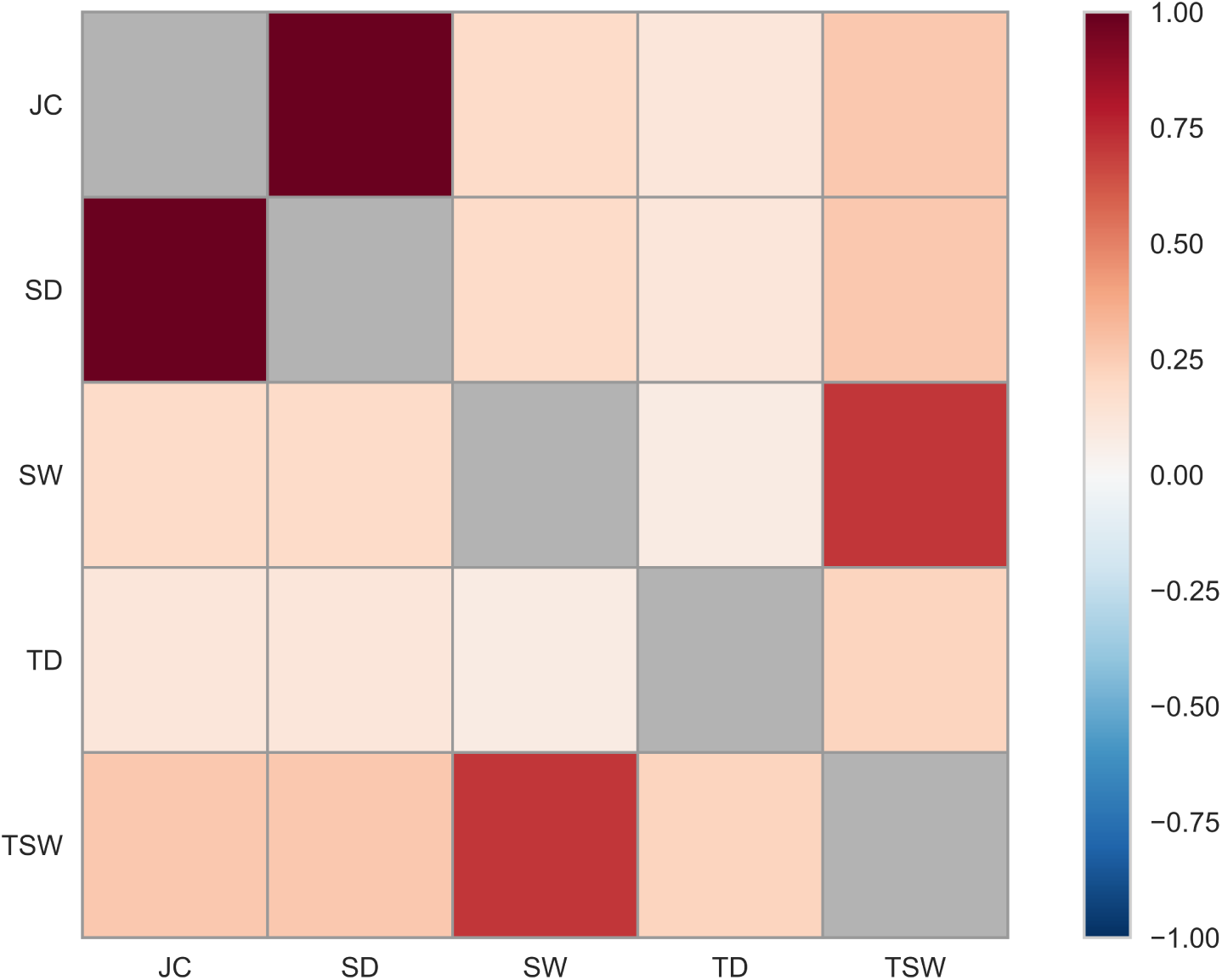
The degree of similarity of functional connectivity estimates for all tested DFC methods computed with the Spearman correlation coefficient in Simulation 1. its covariance over time and how this change must be known beforehand. The aim of this simulation was to see how well the derived DFC estimates can infer the covariance that the data was sampled from when the covariance is fluctuating.

Two time series were generated (*X*). Each time point *t* is sampled from a multivariate Gaussian distribution:

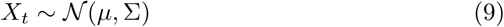

where the covariance matrix was defined as:

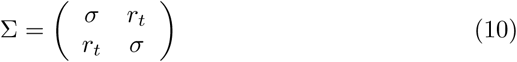

and where the variance, *σ* = 1, was set to 1. At each time point, *r*_*t*_ was sampled from another Gaussian distribution:

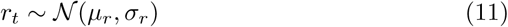

The mean of the time series (*µ*) was set to 0, the mean of the covariance (*µ*_*r*_) was set to 0.2. The variance of the fluctuating covariance (σ_*r*_) was set to vary with the values of {0.08, 0.1, 0.12}.

The covariance at time (*r*_*t*_) was sampled from a Gaussian distribution. Each time point received a new value of *r*_*t*_. This allowed us to compare each DFC method’s connectivity estimate in relation to the time varying covariance parameter *r*_*t*_. Note, *r*_*t*_ is not a definitive measure of the covariance at *t*, it is only a parameter of the covariance used to sample from a Gaussian distribution. Thus, we should not expect the connectivity estimate from any method to correlate perfectly with *r*_*t*_. However, it is possible to compare which method correlate best with *r*_*t*_ to evaluate the overall performance.

The above model will have a temporally fluctuating covariance. It fails to include any autocorrelation in the time series. Not accounting for this may bias the results for some of the tested methods that utilize nearby temporal points to assist estimating the covariance. Merely adding an autocorrelation, like in Simulation 1, will also increase the covariance between the two time series and this will not be tracked by *r*_*t*_. To account for this, we placed a 1-lag autoregressive model for the fluctuating covariance at *r*_*t*_:

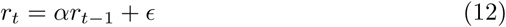

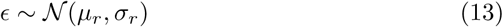

Where α is the autocorrelation parameter. The values for *µ*_*r*_ and σ_*r*_ were the same as above. When *t* = 1, *E* was set to 0.

This revised formulation of our simulation model allowed for the covariance to fluctuate, but with an added autocorrelation on the covariance parameter. In simulation 2, three different settings of the parameter α were used (α =

0, 0.25, 0.5). When α = 0 it is equivalent to the original model outlined above with no autocorrelation. With an increased α it entails a greater influence of the covariance from *t* − 1 in sampling the covariance at *t*. α = 0.5 is reasonable given highly correlated BOLD time series. An α = 0 is more to be expected when time series are less correlated. 10,000 time points were sampled for each of the three different settings of the autocorrelation parameter. See also the Supplementary Materials for a justification of the parameter settings chosen here.

Simulation 2 was run with 9 different simulation hyperparameter combinations: three different values of α and three different values of σ_*r*_.

A sample of time series generated with the model using different settings for the autocorrelation parameter α is shown in Figure 5A. Due to the varying degree of autocorrelation, the mean covariance for time series changes as a function of α, but all still create a Gaussian distribution of *r*_*t*_ (Figures 5B-D). The degree of crosscorrelation between the two time series followed the specified α parameter for the autocorrelation of the covariances (Figures 5E-G).

The results from Simulation 2 are shown in Tables 1-9. The JC method had the lowest WAIC score for all settings of α, followed by the SD method. The SW, TSW and TD methods performed nearly equally well for α = 0. The TD method came in third place for all but one parameter configuration. All WAIC values, their standard error and ∆ WAIC scores are shown in Tables 1-9.

**Table 1:** Results of Simulation 2 where α = 0.0 and σ_*r*_ = 0.08. Tables shows WAIC, WAIC standard error, and difference in WAIC from the best performing method. A lower WAIC indicates a better fit.

| Model | WAIC | WAIC SE | $\Delta$ WAIC |
| --- | --- | --- | --- |
| JC | 28149.6 | 142.529 | 0 |
| SD | 28151.5 | 142.499 | 1.89794 |
| TD | 28206.2 | 142.673 | 56.6558 |
| SW | 28207.6 | 142.662 | 58.0093 |
| TSW | 28207.9 | 142.679 | 58.2634 |

**Table 2:** Results of Simulation 2 where α = 0.0 and σ_*r*_ = 0.1. Tables shows WAIC, WAIC standard error, and difference in WAIC from the best performing method. A lower WAIC indicates a better fit.

| Model | WAIC | WAIC SE | $\Delta$ WAIC |
| --- | --- | --- | --- |
| JC | 28103.6 | 142.963 | 0 |
| SD | 28104.3 | 143.047 | 0.687371 |
| TD | 28200.4 | 143.872 | 96.7158 |
| TSW | 28201.9 | 143.896 | 98.2981 |
| SW | 28205.8 | 143.956 | 102.192 |

**Table 3:**
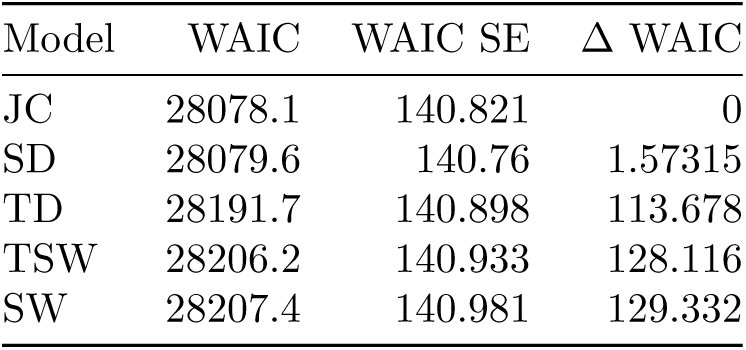
Results of Simulation 2 where α = 0.0 and σ_*r*_ = 0.12. Tables shows WAIC, WAIC standard error, and difference in WAIC from the best performing method. A lower WAIC indicates a better fit.

**Table 4:** Results of Simulation 2 where α = 0.25 and σ_*r*_ = 0.08. Tables shows WAIC, WAIC standard error, and difference in WAIC from the best performing method. A lower WAIC indicates a better fit.

| Model | WAIC | WAIC SE | $\Delta$ WAIC |
| --- | --- | --- | --- |
| JC | 28098.3 | 141.602 | 0 |
| SD | 28101.2 | 141.699 | 2.91403 |
| TD | 28189.5 | 141.813 | 91.1754 |
| TSW | 28206.8 | 141.598 | 108.57 |
| SW | 28207.1 | 141.565 | 108.849 |

**Table 5:** Results of Simulation 2 where α = 0.25 and σ_*r*_ = 0.1. Tables shows WAIC, WAIC standard error, and difference in WAIC from the best performing method. A lower WAIC indicates a better fit.

| Model | WAIC | WAIC SE | $\Delta$ WAIC |
| --- | --- | --- | --- |
| JC | 28104.6 | 139.337 | 0 |
| SD | 28117.6 | 139.386 | 12.9322 |
| TD | 28168.1 | 139.483 | 63.5039 |
| TSW | 28181.8 | 139.501 | 77.1804 |
| SW | 28195.7 | 139.752 | 91.0829 |

**Table 6:** Results of Simulation 2 where α = 0.25 and σ_*r*_ = 0.12. Tables shows WAIC, WAIC standard error, and difference in WAIC from the best performing method. A lower WAIC indicates a better fit.

| Model | WAIC | WAIC SE | $\Delta$ WAIC |
| --- | --- | --- | --- |
| JC | 28043.9 | 140.876 | 0 |
| SD | 28050.1 | 140.796 | 6.22696 |
| TD | 28160.6 | 140.532 | 116.743 |
| TSW | 28191.3 | 140.782 | 147.414 |
| SW | 28205.8 | 140.958 | 161.904 |

**Table 7:**
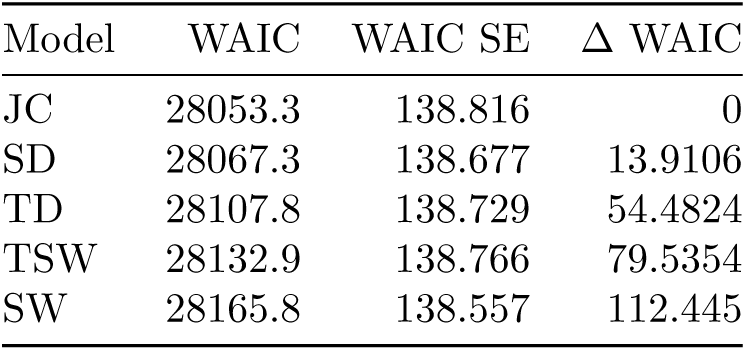
Results of Simulation 2 where α = 0.5 and σ_*r*_ = 0.08. Tables shows WAIC, WAIC standard error, and difference in WAIC from the best performing method. A lower WAIC indicates a better fit.

| Model | WAIC | WAIC SE | $\Delta$ WAIC |
| --- | --- | --- | --- |
| JC | 28053.3 | 138.816 | 0 |
| SD | 28067.3 | 138.677 | 13.9106 |
| TD | 28107.8 | 138.729 | 54.4824 |
| TSW | 28132.9 | 138.766 | 79.5354 |
| SW | 28165.8 | 138.557 | 112.445 |

**Table 8:** Results of Simulation 2 where α = 0.5 and σ_*r*_ = 0.1. Tables shows WAIC, WAIC standard error, and difference in WAIC from the best performing method. A lower WAIC indicates a better fit.

| Model | WAIC | WAIC SE | $\Delta$ WAIC |
| --- | --- | --- | --- |
| JC | 28037.2 | 141.83 | 0 |
| SD | 28053.5 | 141.797 | 16.2988 |
| TD | 28120.2 | 141.398 | 82.9943 |
| TSW | 28148.7 | 141.628 | 111.464 |
| SW | 28201.2 | 142.018 | 163.961 |

**Table 9:** Results of Simulation 2 where α = 0.5 and σ_*r*_ = 0.12. Tables shows WAIC, WAIC standard error, and difference in WAIC from the best performing method. A lower WAIC indicates a better fit.

| Model | WAIC | WAIC SE | $\Delta$ WAIC |
| --- | --- | --- | --- |
| JC | 27933.6 | 142.039 | 0 |
| SD | 27967.3 | 142.002 | 33.7184 |
| TSW | 28042 | 141.938 | 108.408 |
| TD | 28046.3 | 141.97 | 112.708 |
| SW | 28169.3 | 142.095 | 235.72 |

The posterior distribution of the β parameter for each of the DFC methods for all parameter choices are shown in Figure 6. Larger values in the β distribution for a method (i.e. correlating more with *r*_*t*_) conforms with the best fitting models (i.e. lower WAIC score). The SW method performed the worst, followed by the TSW method. The TD method came in third place. SD and JC showed the best performance, with similar posterior distributions of β, although the JC was always slight higher.

There was little difference between the methods when changing the variance of the fluctuating covariance (σ_*r*_). The β values do however scale when σ_*r*_ changes. When σ_*r*_ is smaller, β values decrease.

In sum, the JC method, followed closely by the SD method, showed the best performance in terms of tracking a fluctuating covariance between two time series as performed in Simulation 2. The TD method ranked in third place when there is a higher crosscorrelation between the time series present. The SW and TSW methods showed the worst performance, both in the WAIC score (Tables 1-9) and posterior distributions of β (Figure 6).

**Figure 5:**
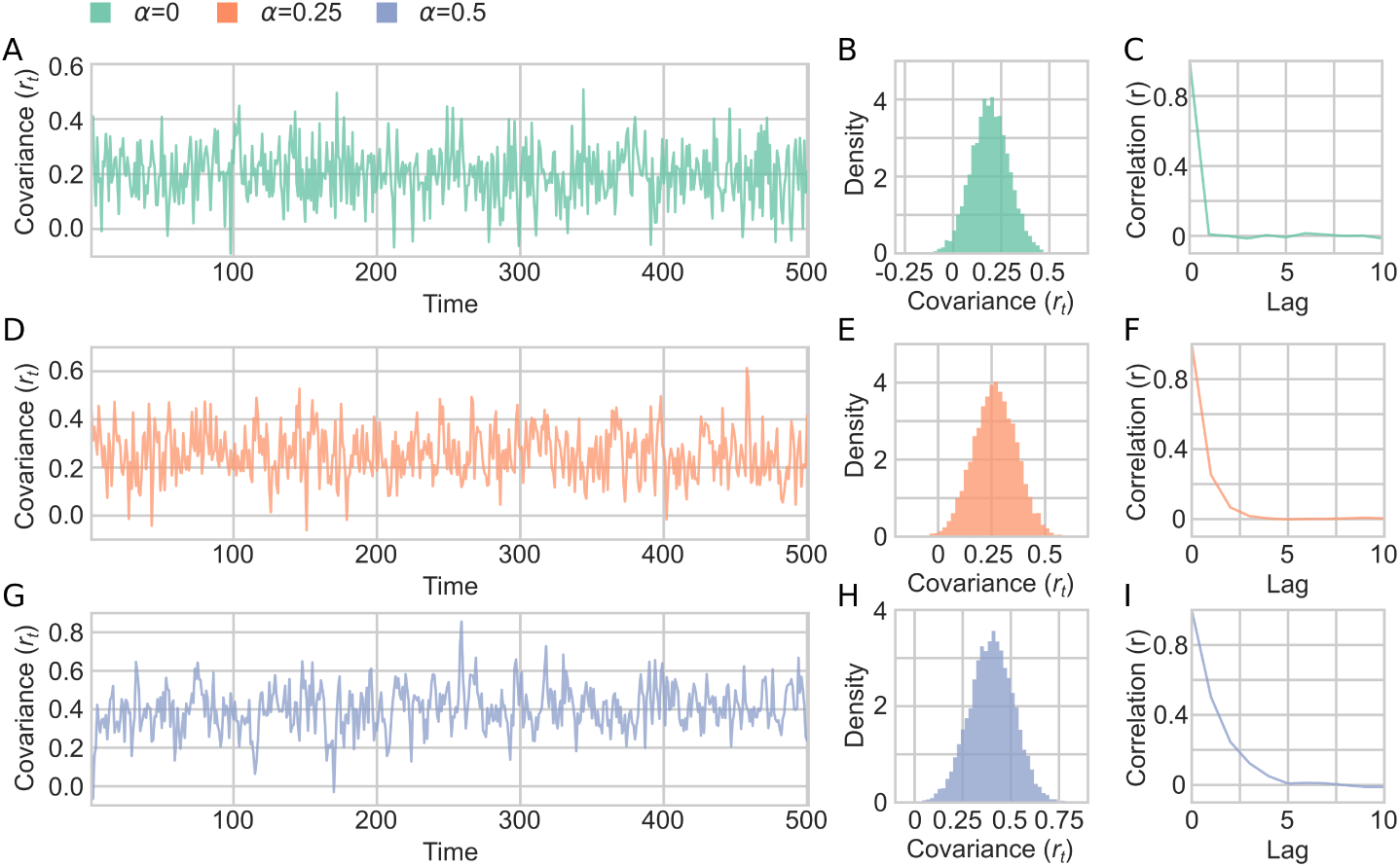
A sample of fluctuating covariance generated in Simulation 2. (A-C) α = 0 and σ_*r*_ = 1. (A) An example of *r*_*t*_ fluctuating over time, showing only first 500 time points shown. (B) Distribution of the fluctuating covariance parameter (*r*_*t*_) (C) autocorrelation of *r*_*t*_ for 10 lags. (D-F) Same as A-C but with the parameters α = 0.25. (G-I) Same as A-C but with the parameters α = 0.5.

**Figure 6:**
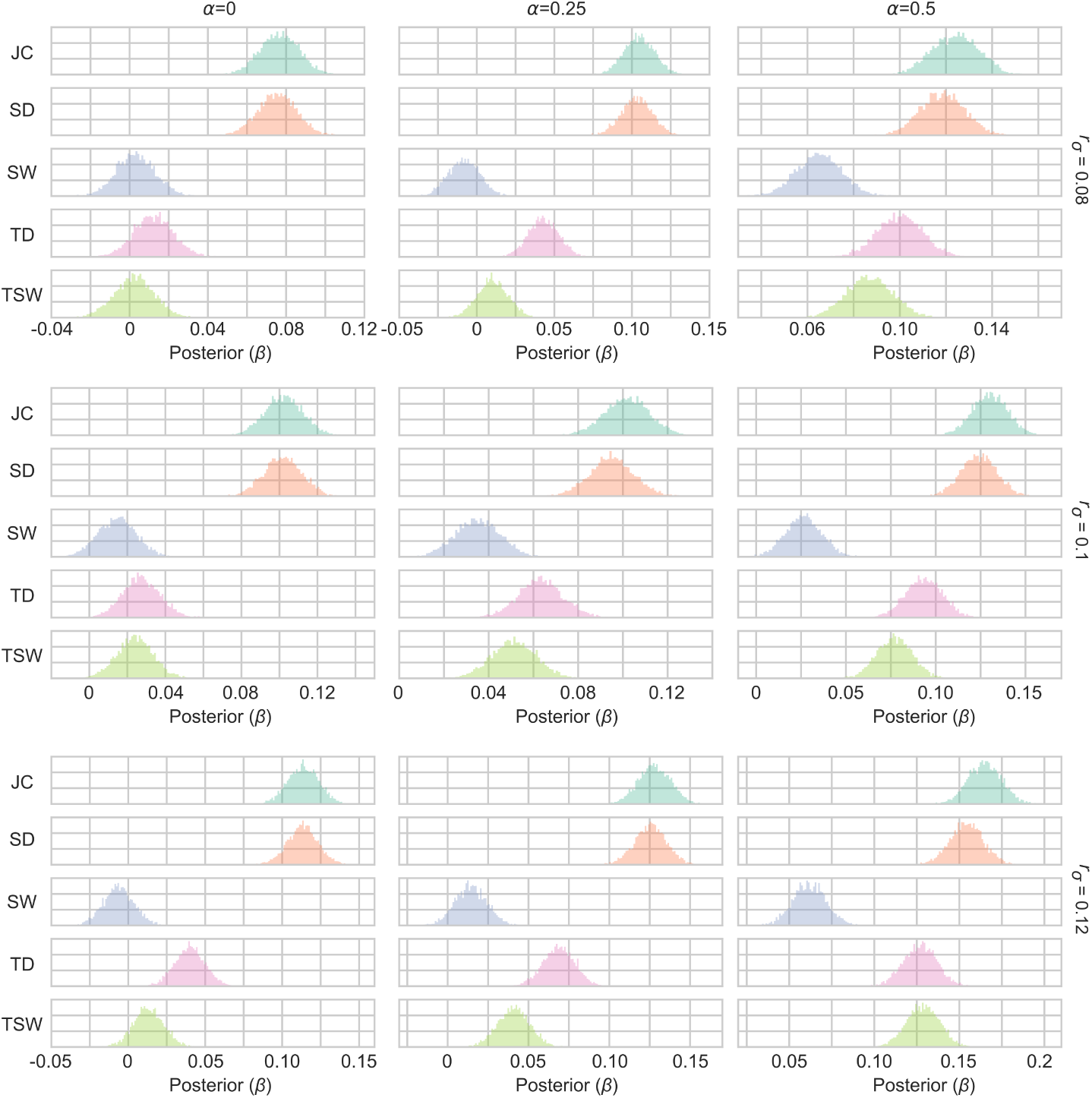
Posterior distributions of the β parameter of the Bayesian linear regression models in Simulation 2. Figure shows a 3x3 grid with the varying values of the autocorrelation (α) and the variance of the fluctuating covariance (σ_*r*_). For each parameter configuration, a model was created for each DFC method. The DFC estimate was the independent variable estimating the fluctuating covariance (*r*_*t*_) between the two time series.

### Simulation 3

The aim of Simulation 3 was to examine the behaviour of different DFC methods when there were non-stationarities present in the data. A typical scenario when this will occur is in DFC is in task fMRI. Simulation 3 is identical in structure to Simulation 2 apart from the following two changes: (1) A non-stationarity, aimed to mimic the occurrence of an event related haemodynamic response function (HRF). Specifically *µ*, which was set to 0 for both time series in Simulation 2, received a different value at each *t* (see next paragraph). (2) σ_*r*_ was set to 0.1 instead of varying across multiple values. This is because Simulation 2 showed no large differences when varying σ_*r*_._*µt*_ was set, for both time series, according to the value of a simulated HRF, that was twenty time points in length and repeated throughout the simulation. The HRF was simulated, with a TR of 2, using the canonical HRF function as implemented in SPM12 using the default parameters (38). This HRF, which has a length of 17 time points, was padded with an additional 3 zeros. The amplitude of the normalized HRF was multiplied by 10 to have a high amplitude fluctuations compared to the rest of the data. *µ*_*t*_ is thus the padded HRF repeated throughout the entire simulated time series. This represents a time series that includes 250 “trials” that each lasts 40 seconds. This simulation helps illustrate how well DFC methods could be implemented in task based fMRI. Examples of the time series generated using different autocorrelation are shown in Figure 7.

The results from Simulation 3 are shown in Figure 8 (Posterior distributions of β) and Tables 10-12 (model fit) which evaluated each DFC’s method performance at tracking the fluctuating covariance (*r*_*t*_). Results were similar with Simulation 2. In the case when the autocorrelation of the covariance was 0, the SW, TSW and TD methods performed quite poorly, but all improved to varying degrees as this increased. The SW method was generally the worst method, followed by TSW. The TD method came in third place. The JC method has the best performance, followed closely by the SD method, in all parameter conditions.

**Table 10:**
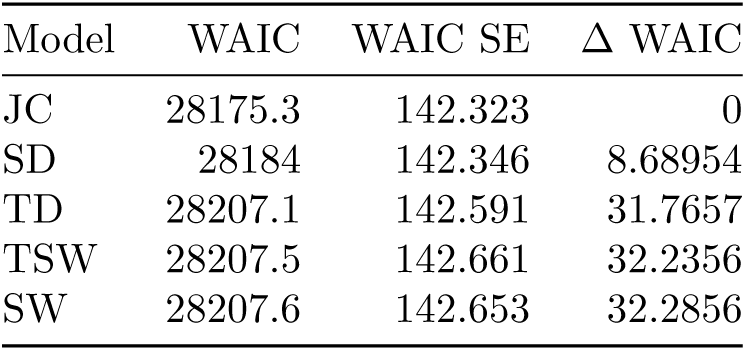
Results of Simulation 3 where α = 0.0. Tables shows WAIC, WAIC standard error, and difference in WAIC from the best performing method. A lower WAIC indicates a better fit.

| Model | WAIC | WAIC SE | $\Delta$ WAIC |
| --- | --- | --- | --- |
| JC | 28175.3 | 142.323 | 0 |
| SD | 28184 | 142.346 | 8.68954 |
| TD | 28207.1 | 142.591 | 31.7657 |
| TSW | 28207.5 | 142.661 | 32.2356 |
| SW | 28207.6 | 142.653 | 32.2856 |

**Table 11:** Results of Simulation 3 where α = 0.25. Tables shows WAIC, WAIC standard error, and difference in WAIC from the best performing method. A lower WAIC indicates a better fit.

| Model | WAIC | WAIC SE | $\Delta$ WAIC |
| --- | --- | --- | --- |
| JC | 28138.3 | 142.606 | 0 |
| SD | 28160.5 | 142.749 | 22.2079 |
| TD | 28184.1 | 143.06 | 45.8161 |
| TSW | 28190.6 | 143.249 | 52.3508 |
| SW | 28202 | 143.265 | 63.7404 |

**Table 12:** Results of Simulation 3 where α = 0.5. Tables shows WAIC, WAIC standard error, and difference in WAIC from the best performing method. A lower WAIC indicates a better fit.

| Model | WAIC | WAIC SE | $\Delta$ WAIC |
| --- | --- | --- | --- |
| JC | 28106.1 | 139.883 | 0 |
| SD | 28136.7 | 139.769 | 30.6289 |
| TD | 28150.5 | 139.587 | 44.4145 |
| TSW | 28177.2 | 139.546 | 71.1701 |
| SW | 28203 | 139.717 | 96.9538 |

In sum, the results from Simulations 2 and 3 suggests that the JC method has the best performance in terms of detecting fluctuations in covariance compared to the other four DFC methods. This result also holds when a non-stationary event related haemodynamic response was added to the mean of the time series.

### Simulation 4

Simulation 4 aimed to test how sensitive different DFC methods are to large and sudden changes in covariance (i.e. changes in “brain state”) that previously have been postulated to exist in fMRI data (e.g. (11,15,17)). We here start in a similar fashion as we did in Simulation 2 where samples for the two time series are drawn from a multivariate Gaussian distribution

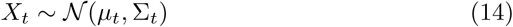

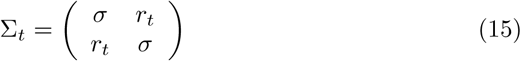

**Figure 7:**
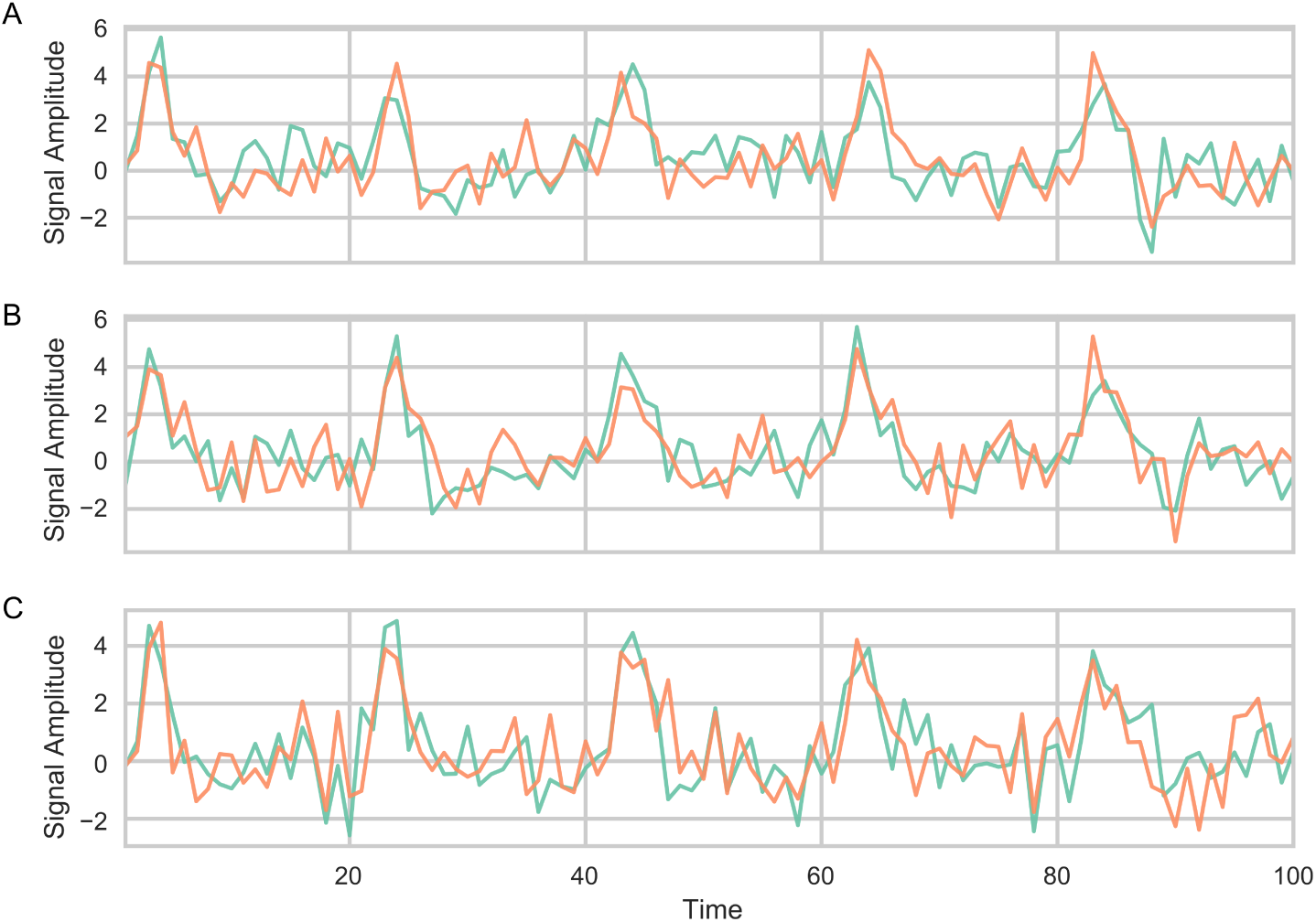
Examples of time series used in Simulation 3 where the mean of the time series is sampled from a time series that included a train of simulated event related HRF fMRI responses (spaced apart every 20 time points). Only the first 100 time points are shown. (A) α = 0, (B) α = 0.25, (C) α = 0.5

**Figure 8:**
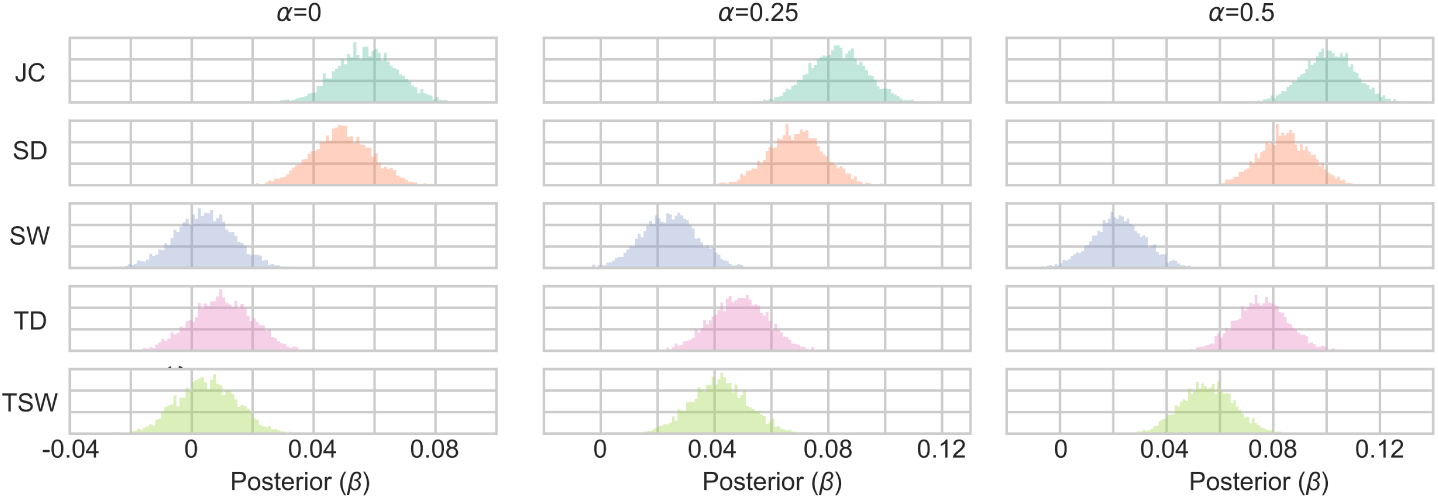
Posterior distributions of the β parameter of the Bayesian linear regression models in Simulation 3. Figure shows a 1x3 grid with the varying values of the autocorrelation (α). For each parameter configuration, a model was created for each DFC method. The DFC estimate was the independent variable estimating the fluctuating covariance (*r*_*t*_) between the two time series.

Similar to simulation 2, we set *µ*_*t*_ = 0 and σ = 1. The covariance parameter *r*_*t*_ was sampled from a Gaussian distribution where the mean was shifted

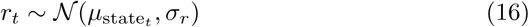

and where σ_*r*_ = 1. At each state transition, *µ*_state_ was randomly chosen from a set *M* (*M* = 0.2, 0.6). The duration of each state was randomly sampled from *L*. Two different scenarios for state transitions were simulated. In the fast transition condition *L* = *{*2, 3, 4, 5, 6*}* and in the slow transition condition

*L* = *{*20, 30, 40, 50, 60*}*. These values correspond to the number of time points a “state” lasts. Beginning at *t* = 1, *µ*_state_ ^to *µ*^state

*t*+*l* was randomly sampled from *M* where *l* was sampled from *L*. This procedure was continued until *X*_*t*_ was 10,000 samples long.

These choices for brain state changes provide time scales of state transitions between 40-120 seconds (slow condition) or 4-12 seconds (fast condition) in simulated fMRI data with a TR of 2 (Figure 9A). The statistical model for evaluating the different DFC methods performance was the same as Simulation 2 and 3. A summary of data generated in Simulation 4 is shown in Figure 9A-F.

The results from Simulation 4 are shown in Figure 10 and Tables 13-14. In the quick transition condition, the JC and the SD showed the best performance for both the WAIC scores and the posterior distribution of β (Figure 10A; Table 7). In the slow transition condition the two sliding window methods outperformed the other methods (Figure 10B; Table 8). The JC and SD methods perform similarly for both conditions. Thus, when there are shifts in covariance that occur relatively slowly, the sliding window methods are sensitive at tracking these changes.

**Table 13:** Results of Simulation 4 where State length = {2,3,4,5,6}. Tables shows WAIC, WAIC standard error, and difference in WAIC from the best performing method. A lower WAIC indicates a better fit.

| Model | WAIC | WAIC SE | $\Delta$ WAIC |
| --- | --- | --- | --- |
| JC | 27548.3 | 92.0207 | 0 |
| SD | 27571.1 | 92.7548 | 22.8124 |
| TD | 27741.6 | 93.5845 | 193.243 |
| TSW | 27749.5 | 93.3275 | 201.18 |
| SW | 28072.5 | 87.9986 | 524.197 |

**Table 14:**
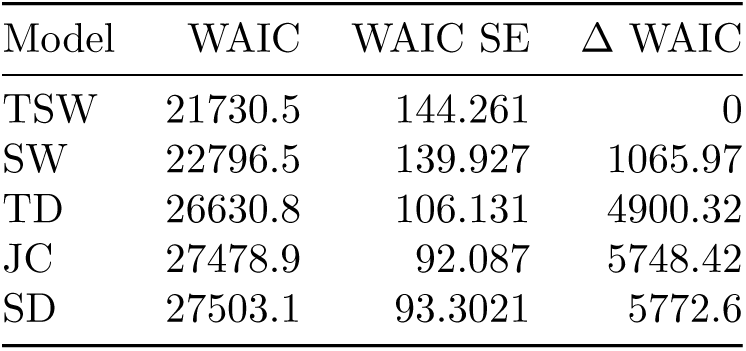
Results of Simulation 4 where State length = {20,30,40,50,60}. Tables shows WAIC, WAIC standard error, and difference in WAIC from the best performing method. A lower WAIC indicates a better fit.

## Discussion

In this study we have developed four simulations to test the performance of different proposed dynamic functional connectivity methods. The first simulation showed which methods yield similar connectivity time series. Notably, all methods correlated positively with each other, but to a varying degree. The second simulation generated data in which the autocorrelated covariance between simulated time series varied in time. In this case, the JC method, followed closely by the SD method, showed the best performance. In the third simulation, the generated time series contained a non-stationary mean related to haemodynamic responses. Again, our simulations suggested that the JC method performed best. The forth simulation included nonlinear shifts in covariance (in an attempt to simulate brain state shifts). When the states changes were quick, the JC method performed best. When the state changes were slow, the TSW (followed by the SW) performed best.

**Figure 9:**
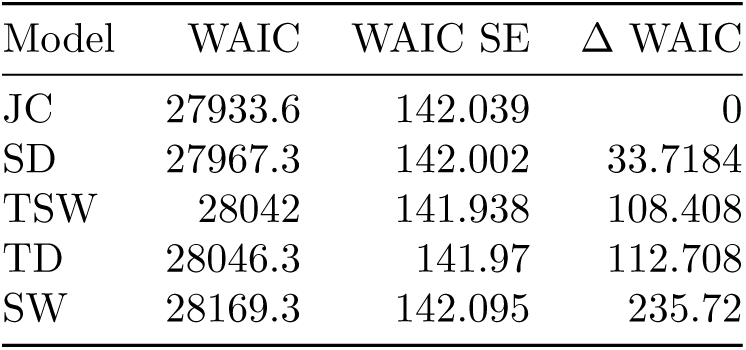
A sample of fluctuating covariance generated in Simulation 4. (A-C) Quick state transitions (between 2-6 time points long). (A) An example of *r*_*t*_ fluctuating over time, showing only first 500 time points shown. (B) Distribution of the fluctuating covariance parameter (*r*_*t*_) (C) autocorrelation of *r*_*t*_ for 10 lags. (D-F) Same as A-C but with the long state transitions (between 20-60 time points long).

**Figure 10:**
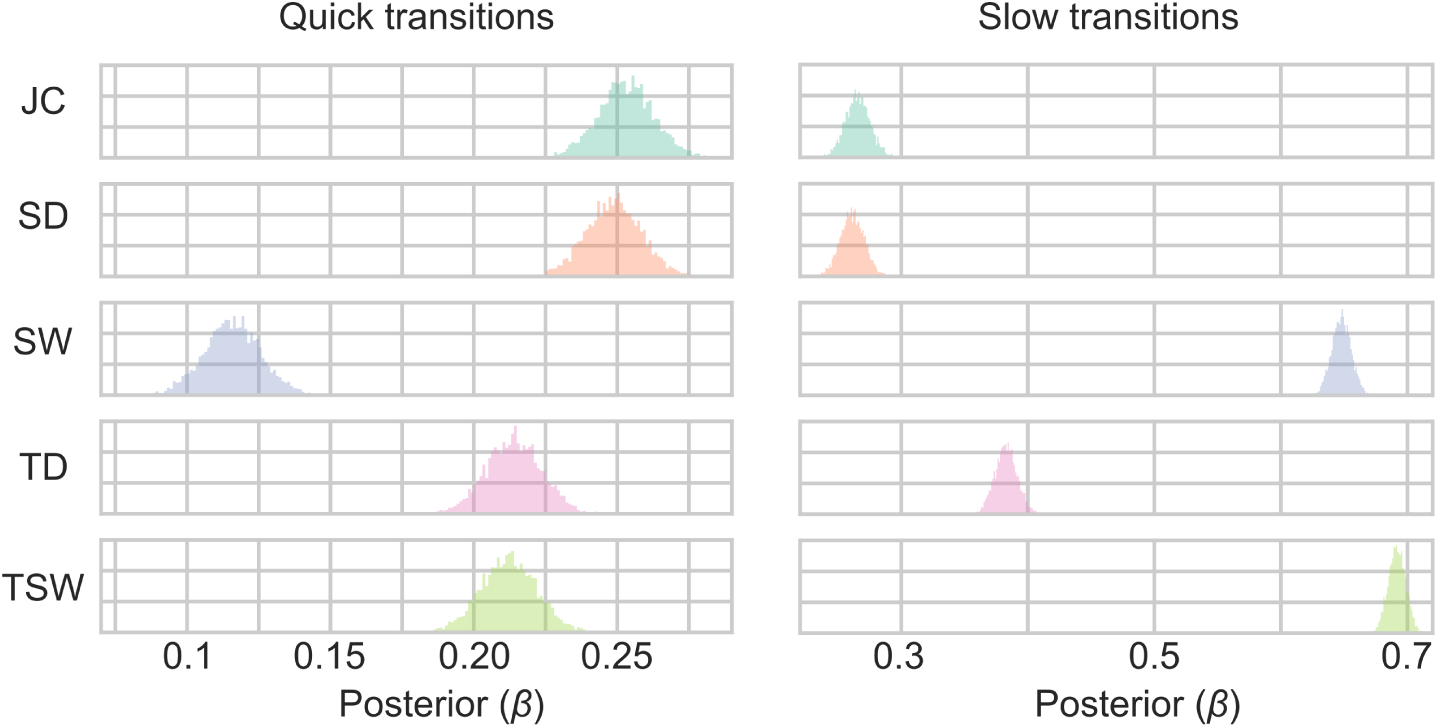
Posterior distributions of the β parameter of the Bayesian linear regression models in Simulation 4. Figure shows a 1x2 grid with the varying values of the state length. For each parameter configuration, a model was created for each DFC method. The DFC estimate was the independent variable estimating the fluctuating covariance (*r*_*t*_) between the two time series.

In a previous simulation that evaluated the sliding window method, the sensitivity of SW and TSW method was found to be good at detecting state shifts (39). Here, at least when the transitions are slow, we found similar results. The sliding window methods is optimal if there are slow state changes. However it is unclear if “state changes” are the best yardstick for dynamic brain connectivity. In particular, non-stationarities in dynamic connectivity have been attributed to spurious sources such as movement (12). Given the unknowns of the “true” connectivity, methods which are robust over conditions are more likely the safer options — in this case the JC or SD method performed similarly in both conditions.

Overall the jackknife correlation method performed the best across all simulations. We have shown it to be robust to numerous changes in parameters. However, the JC method is not without some considerations. First, it introduces variance compression that reduces the absolute variance, while preserving the relative variance within the time series. This variance compression also scales with the length of the time series. The consequence of this is that direct comparisons of the DFC variance between cohorts/conditions becomes hard to interpret as dynamic fluctuations, especially when the length of the data varies. However, this is the case for most methods and it should be remembered that the variance is proportional to the static functional connectivity (7,9,10). Simply put, the JC method (like all other methods) should not be used for a direct contrast of the variance of DFC time series. Second, the JC method sensitivity means that noise will be carried over per time point instead of being smeared out over multiple time points. This is actually beneficial as it allows for further processing steps to be applied that aims to remove any remaining noise (e.g. motion) which cannot be done when the noise has been smeared across the connectivity time series (e.g. in windowed methods).

The simulations and results presented in this study should not be taken as an exhaustive and complete assessment of all aspects of a given method to conduct DFC. Rather, the four simulations described here represents a subset of possible scenarios in terms of different methodological characteristics that might be of interest. The current four simulations are marked *dfcbenchmarker* simulation routine V1.0. If modifications or additional scenarios are considered to be improvements to the current simulations, these will get an updated version number. Many additional simulations could be conceived of on top of this original routine. For example, one could include multiple time series, adding movement type artifacts, adding frequency relevant characteristics, a stationary global signal etc. These have not been included here, as the focus in these simulations was to primarily assess tracking of a fluctuating covariance. Input from researchers about appropriate additions to the simulations is welcome.

We encourage researchers designing DFC methods to benchmark their own results with *dfcbenchmarker* (www.github.com/wiheto/dfcbenchmarker). Researchers need only to write a Python function for their method and use it as an input for dfcbenchmarker.run_simulations() and their method will be compared to the five methods presented in this paper (see online documentation). Functions can then be submitted through the function dfcbenchmarker.send_method(). All valid methods submitted will be released in summaries of the submitted benchmarked results so that researchers can contrast the performance of different methodologies.

## Author contributions

WHT and PF designed the simulation analysis. WHT and CGR designed and tested the jackknife correlation method. WHT and PPS designed the statistical models. WHT performed the analysis. WHT and PF wrote the paper. CGR and PPS critically revised the paper.

## Acknowledgements

We would like to thank Russell Poldrack and Lieke de Boer for helpful comments and discussions about the manuscript. This work was supported by the Swedish Research Council (grants no. 2016-03352 and 773 013-61X-08276-26-4) and the Swedish e-Science Research Center.

## Supplementary Materials - A justification of the assumptions regarding model parameter choices made in the simulations

In simulation routine V1.0 of dfcbenchmarker, we wanted the simulations to either be justified based on the properties of fMRI data or, in the case of Simulation 4, reflect what dynamics are hypothesized to exist in fMRI data.

Since the properties of the simulations are determined by their parameters, we justify these parameters in three different ways. First, we show that some parameters have no substantial effect on the result. Second, some are justified from previous findings/hypothesis in the literature. Third, by use resting state fMRI data to justify the assumptions.

### fMRI data used

The midnight scanning club (MSC) data (1) is used to justify the assumptions. The dataset contains ten subjects with ten sessions of resting state fMRI. This data was obtained from the OpenfMRI database (accession number: ds000224). The preprocessed volumetric data was used. To reduce the dimensions of the data, 278 spherical regions of interest were defined (5mm radius) with their the centre of mass defined by the Shen et al 2013 atlas (2) (using the Talairach coordinates). Subject MSC08 is often reported as an outlier, which has previously been noted in Gordon et al where they noted this subject had poor data quality due to subject head movement and sleep (1). When a single subject and and session is shown (for illustration purposes), subject MSC10 and session 7 was used. This subject and session was selected at random. When specific ROIs were selected to be shown, these were randomly generated (from subject MSC10, session 7).

See github.com/wiheto/dfcbenchmarker_assumptions/1.0/assumptions.ipynb for the Jupyter notebook to run all the analysis from start to finish.

### Simulation 1 - summary of assumptions

#### Information

*Aim*: See how well the estimates from different DFC methods correlate with each other.

*Evaluation*: Spearman correlation between functional connectivity time series computed by the five DFC methods.

*n samples*: 10,000

*Random seed*: 2017

#### Assumptions

*Distributions*: Gaussian, multivariate Gaussian

**µ* (mean of each time series)*: 0

*o (variance of time series)*: 1

*r_t_ (covariance of time series)*: 0.5

*α (autocorrelation of the time series)*: 0.8

### Simulation 2 - summary of assumptions

#### Information

*Aim*: To investigate how well the different DFC methods correlate with a fluctuating covariance parameter.

*Evaluation*: Bayesian linear regressions between the estimates of the DFC methods and *r*_*t*_ (evaluated by comparing WAIC scores and posteriors of β).

*n samples*: 10,000

*Random seed*: 2017

#### Assumptions

*Distributions*: Gaussian, multivariate Gaussian

**µ* (mean of each time series)*: 0

*o (variance of time series)*: 1

_*µr*_ (mean of fluctuating covariance): 0.2

_*σr*_ (variance of fluctuating covariance): {0.08, 0.1, 0.125}

*α (autocorrelation of the fluctuating covariance)*: {0, 0.25, 0.5}

### Simulation 3 - summary of assumptions

#### Information

*Aim*: To investigate how well different DFC methods perform when the fluctuating covariance parameter contains a non-stationary mean to simulate a HRF.

*Evaluation*: Bayesian linear regressions between the estimates of the DFC methods and *r*_*t*_ (evaluated by comparing WAIC scores and posteriors of β).

*n samples*: 10,000

*Random seed*: 2017

*TR*: 2

#### Assumptions

*Distributions*: Gaussian, multivariate Gaussian

*µ* A scaled HRF.

*o (variance of time series)*: 1

_*µr*_ (mean of fluctuating covariance): 0.2

_*σr*_ (variance of fluctuating covariance): 0.1

*α (autocorrelation of the fluctuating covariance)*: {0, 0.25, 0.5}

*Trial length (samples)*: 20

*HRF scale*: 10

### Simulation 4 - summary of assumptions

#### Information

*Aim*: To investigate the performance of DFC methods when there is a fluctuating covariance parameter that nonlinearly shifts between covariance.

*Evaluation*: Bayesian linear regressions between the estimates of the DFC methods and *r*_*t*_ (evaluated by comparing WAIC scores and posteriors of β).

*n samples*: 10,000

*Random seed*: 2017

#### Assumptions

*Distributions*: Gaussian, multivariate Gaussian

*µ (mean of each time series)*: 0

*o (variance of time series)*: 1

*rµ (average covariance in different states)*: 0.2, 0.6

*rσ (variance of covariance in different states)*: 0.1

*State length*: Fast Condition: {2,3,4,5,6}; Slow condition: {20,30,40,50,60}

#### Justifying the assumptions

##### Mean and variance of the time series (*µ*, σ)

*µ* and σ for all simulations were set to 0 and 1, with the exception of *µ* in simulation 3. As long as the time series have a stationary mean and variance, then these parameters can be set to anything and have no effect on the overall result. Simulation 3 tests how methods deal with the non-stationartity of *µ*.

##### Autocorrelation and crosscorrelation assumptions (α) Autocorrelation of the time series (Simulation 1)

The auto-correlation (α) is an important parameter for simulation 1. In Figure S1, we can see the average auto-correlation for a single subject between 0 and 10 lags. The average and standard deviation for each subjects and session is shown in Figure S2. Excluding the sessions from the subject known to be noisy (MSC08), there were only a few sessions with average auto-correlation values outside the range of 0.70-0.85. Most subject averages were close to 0.80. Accordingly, α in simulation 1 was set to 0.80.

##### Autocorrelation of *r*_t_ (Simulation 2 and 3)

The autocorrelation of *r*_*t*_ is an important parameter for simulation 2 and 3, where the parameter α now refers to the autocorrelation of the covariance parameter *r*_*t*_. The expected crosscorrelation will be different for different values of static functional connectivity. Thus, we binned each edge based on its static connectivity bins (each bin was placed between -1 and 1 in spaces of 0.1). Figure S3A shows the average crosscorrelation of the example subject and session for 10 lags for each bin. The general pattern is that at lag 1, the average correlation tends towards zero compared to lag 0 (which is the static functional connectivity). The frequency of the number of edges across the different correlation values is at lag 0 (Figure S3B) and lag 1 (Figure S3C) shows that the majority of edges are within the range -0.5 and 0.5.

The average crosscorrelation at lag 1 looks very similar across different subjects and sessions (Figure S4). When pooling all static functional connectivity values across subjects and sessions, the histogram (Figure S5) shows that the pattern at a group level is similar to that shown in Figure S3B. Thus, more edges correspond to a degree of connectivity that would be modeled with a lower α parameter. However, for edges with larger degree functional connectivity, a higher α is to be expected.

**Figure 1:**
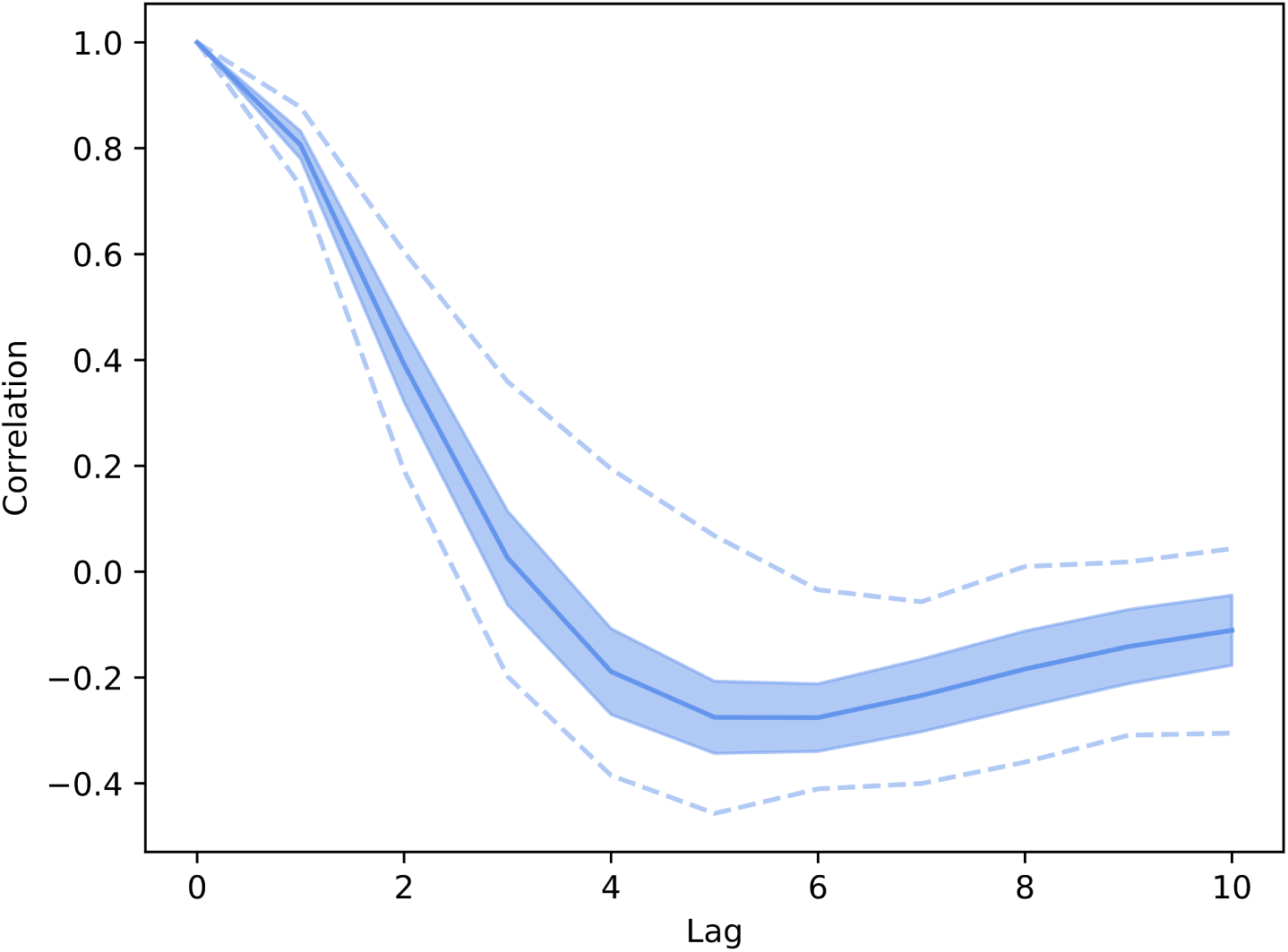
Auto-correlation of the example subject and session for 10 lags. Averaged over all ROIs. Error bars show standard deviation. Dashed lines show the min and max.

**Figure 2:**
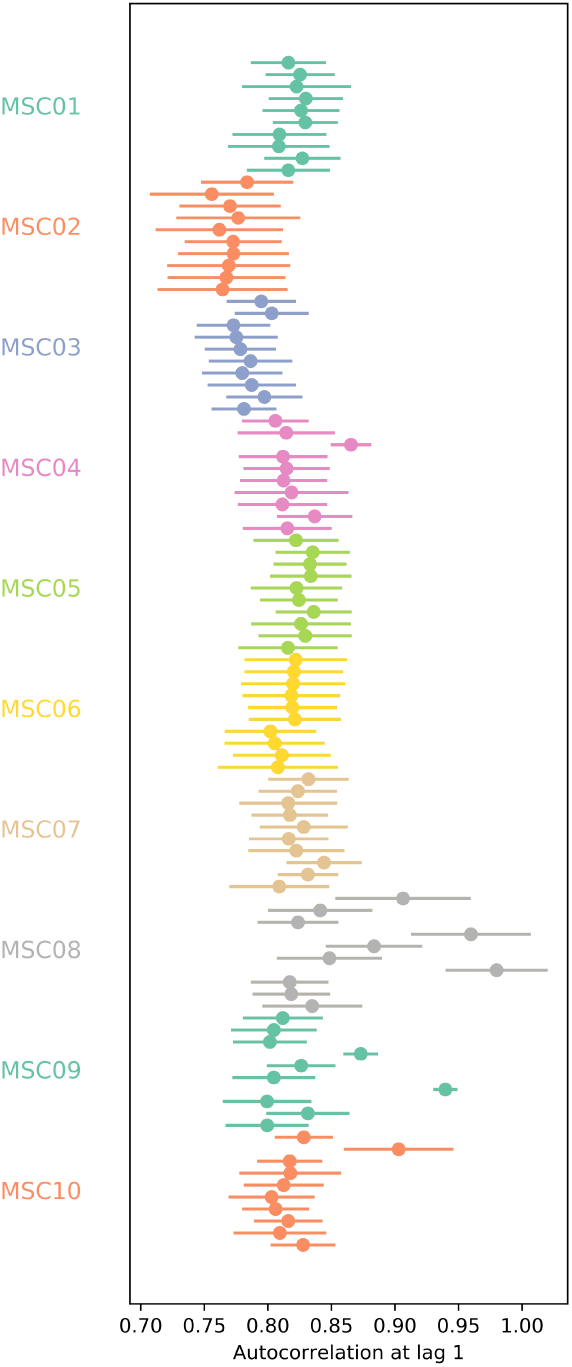
Average auto-correlation at lag 1 for each subjects and sessions. Each dot represents a session (ordered from top to bottom). Each dot signifies the average autocorrelation at lag 1. Error bars show standard deviation.

**Figure 3:**
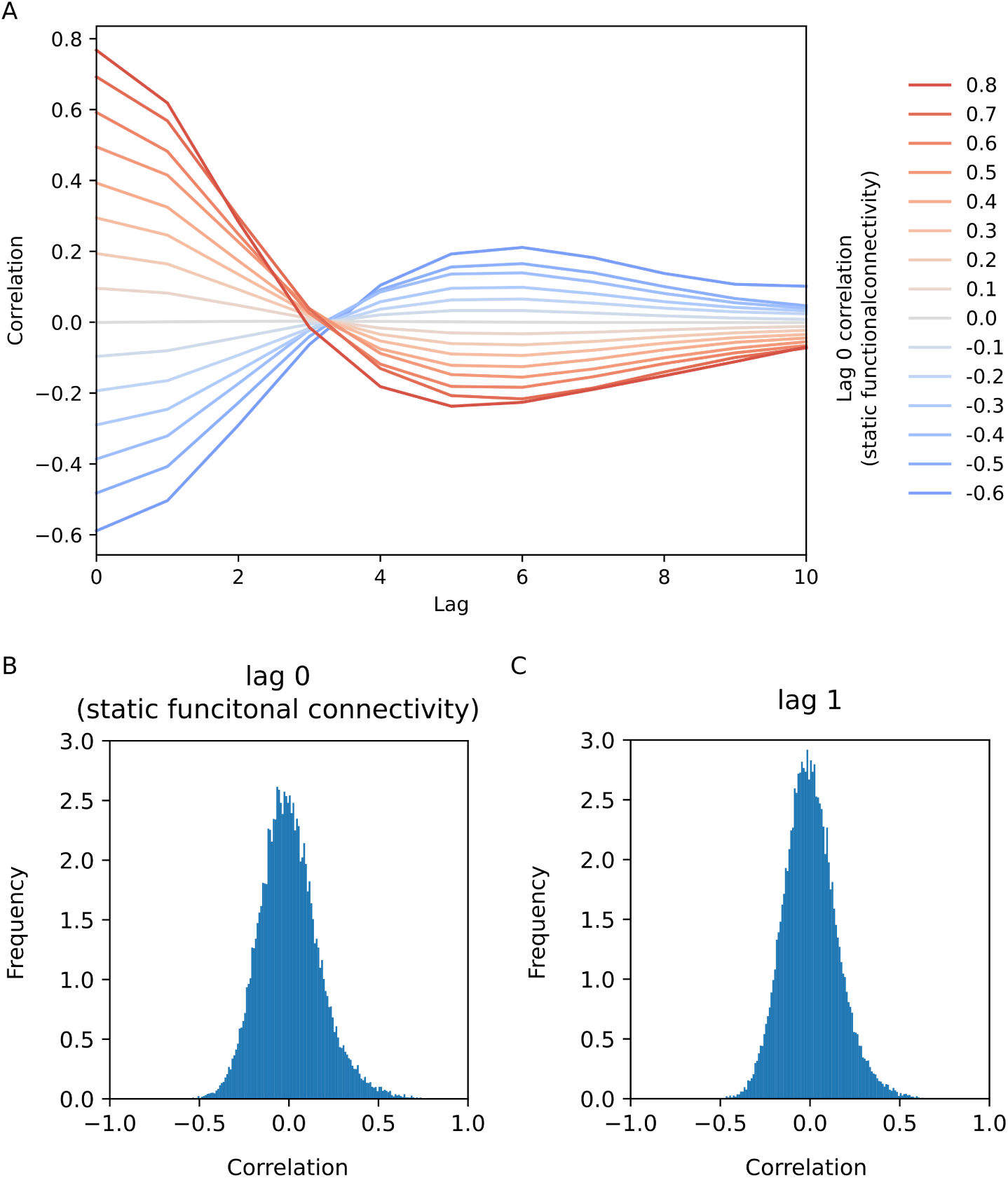
Average crosscorrelation of the example subject and session for 10 lags. Averaged over all ROIs. Each coloured line represents a bin based on the correlation value at lag 0 (i.e. the static functional connectivity).

**Figure 4:**
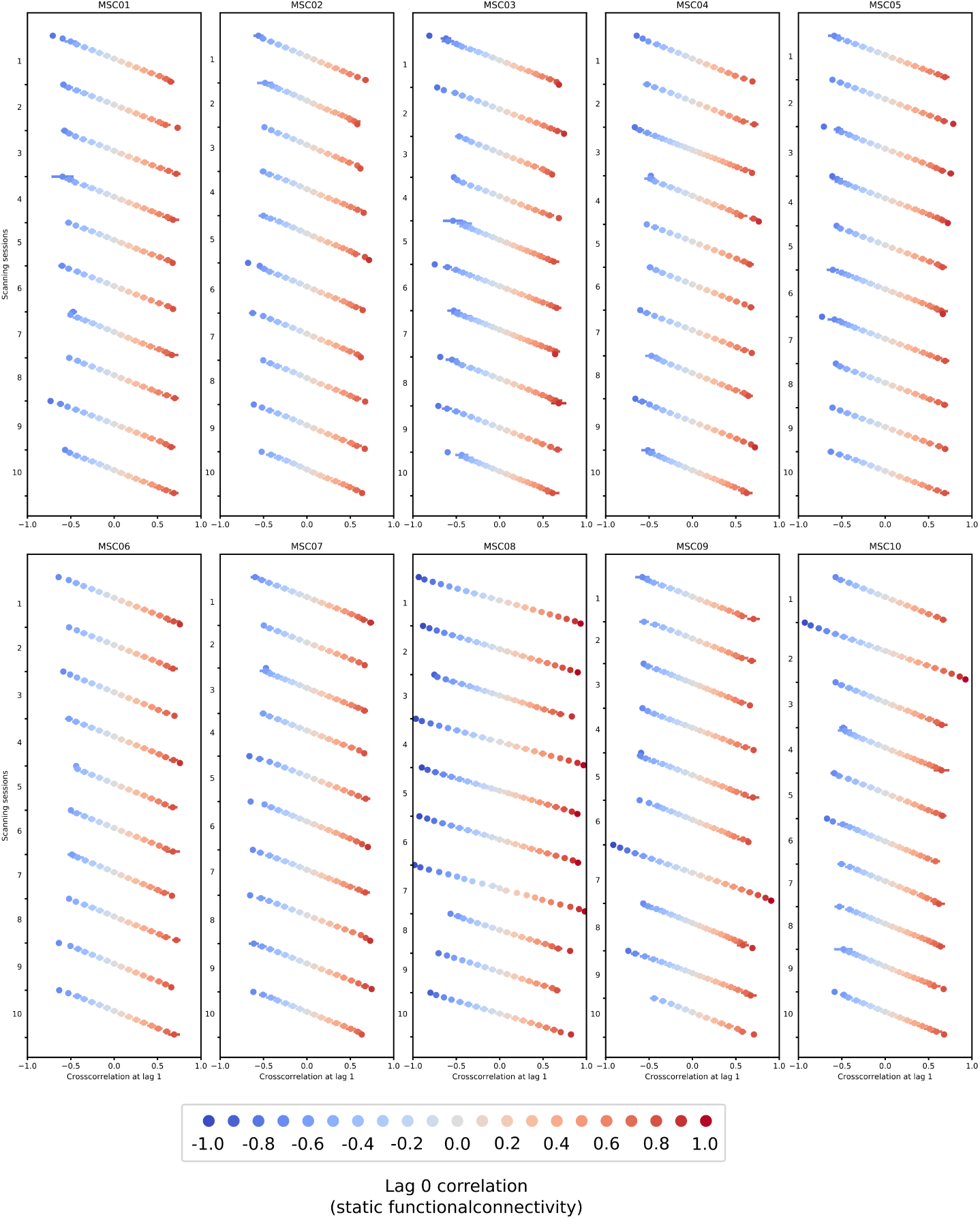
Average crosscorrelation at lag 1 for all subjects and sessions. Each dot represent a bin based on the correlation value at lag 0 (i.e. static functional connectivity). Each dot signifies the average (across ROIs in the bin) crosscorrelation at lag 1. Error bars indicate standard deviation.

**Figure 5:**
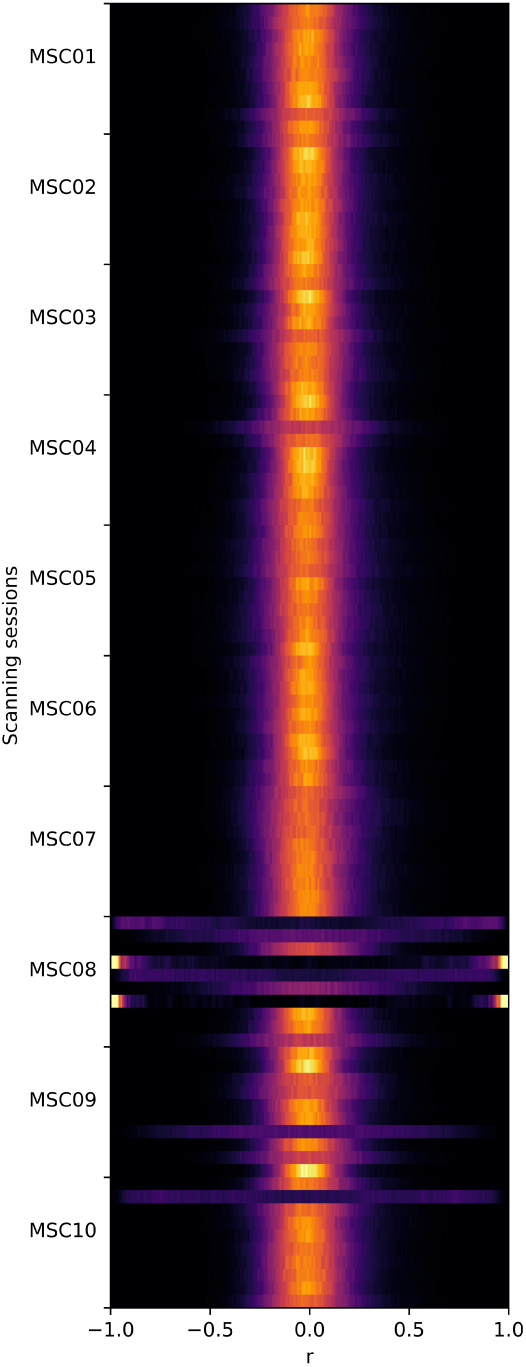
The frequency of the static functional connectivity for each subject and session. Each row in the figure represents one session for a subject. Each row contains a normalized histogram of the connectivity values.

When choosing the α for simulation 2 and 3, we only considered positive auto-correlations and sampled from the values 0, 0.25, and 0.5 which cover most of the positive span of functional connectivity.

##### Covariance (*µ*_*r*_, σ_*r*_, *r*_*t*_)

_*σr*_ dictates how wide *r*_*t*_ is going to fluctuate over time. What the true value of σ_*r*_ is, is one of the key questions dynamic functional connectivity wants to answer. Making σ_*r*_ higher, entails that there is a larger parameter space that *r*_*t*_ gets sampled from which decreases the noise when estimating *r*_*t*_. When the distribution that *r*_*t*_ is sampled from widens, there is a lower probability that the time series will be sampled similarly, meaning the inherent uncertainty of estimating single samples of *r*_*t*_ decreases when the variance of *r*_*t*_ increases. The effect of changing σ_*r*_ is that the β values will scale with it, but the relative difference of the β values between methods will remain similar. The σ_*r*_ parameter in is set to 0.08, 0.1 and 0.12 to illustrate this in simulation 2 to show that this has little effect other than a scaling effect of β (Figure 6). In simulation 3, only 0.1 was used for σ_*r*_.

The average connectivity (*µ*_*r*_) was set to 0.2 in Simulation 2 and 3. It can be seen in Figure S5 that this is a reasonable positive connectivity value. The *µ*_*r*_ parameter is unchanged throughout simulation 2 and 3, however due to the fluctuating α parameter the effective covariance between the two time series increases as a function of α (Figure 5). It can be seen that this effective increase in covariance had little effect on the different methods (Figure 6, Figure 8).

The *r*_*t*_ in simulation 4 switched between a very high connectivity “state” of 0.6 and lower, but positive, connectivity “state” of 0.2. These are plausible connectivity values (Figure S5). See the section below on state change assumptions for more details.

### Gaussian assumption

All simulated time series were sampled from a (multivariate) Gaussian distribution. The simulations can be improved by considering frequency specific information (see caveats below). The amplitude of the time series for ROIs in the example subject and session have a unimodal shape and relatively unskewed distributions (Figure S6A). To illustrate this more clearly, Figure S6B shows three randomly selected ROIs. When looking at all groups the average skewness was calculated (Figure S7). Given the distributions are unimodal they too can at least be approximated with a Gaussian distribution. While not perfect, and still a simplification, a Gaussian distribution is a reasonable distribution to sample from. The Gaussian distribution assumption for sampling *r*_*t*_ can also be motivated by (3) where the dynamic functional connectivity was generally unimodal and, with appropriate transformation, approximates to Gaussian.

**Figure 6:**
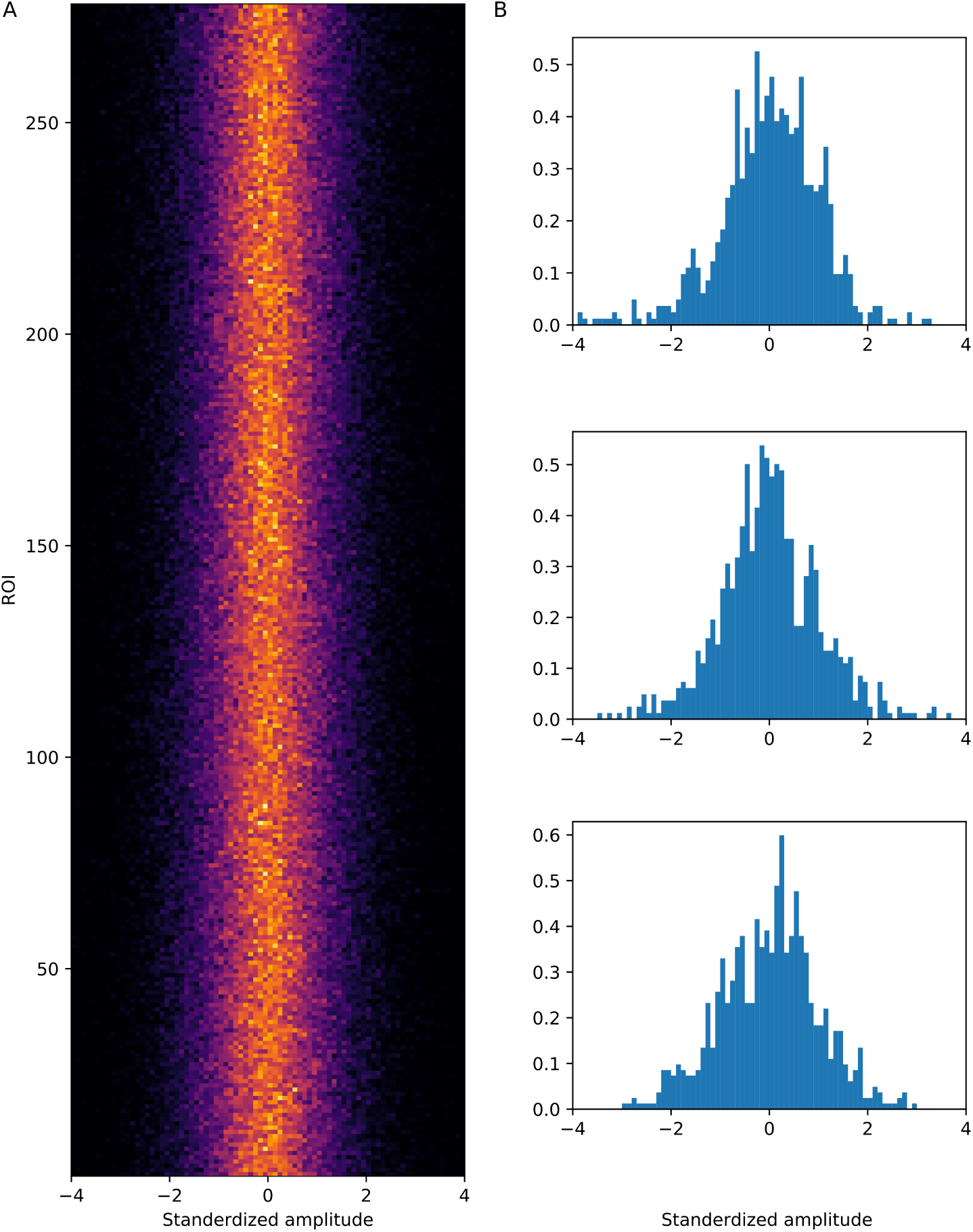
Distribution of BOLD signal for each ROI in example subject and session. A. Histograms over each ROI showing the frequency of signal intensity. Each row functions as a histogram for a ROI. B. Three randomly selected ROIs from A are depicted as traditional histograms showing the same information as in A.

**Figure 7:**
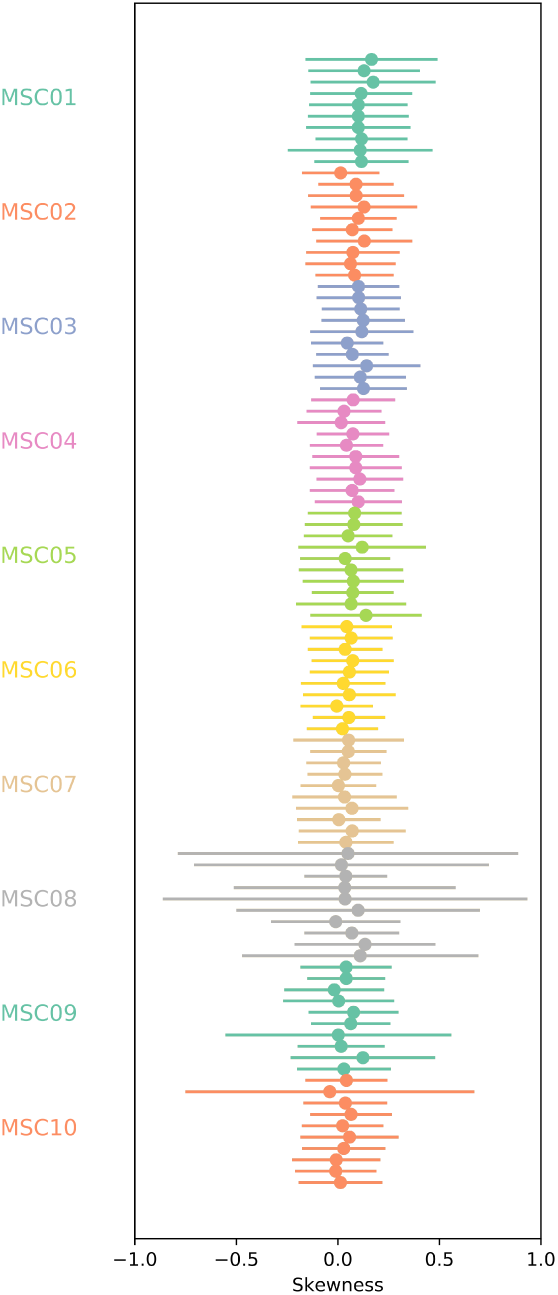
Average skewness for each subject and session. Each dot represents a session. Each dot signifies the average skewness over all BOLD time series for ROIs. Error bars show one standard deviation appropriate transformation, approximates to Gaussian.

### Trial length and HRF scale

The scale of the HRF amounts to how much the standardized HRF has been scaled so that it creates a larger non-stationarity in mean of the simulated data in Simulation 3. This parameter was set to a high value to illustrate each method’s ability to estimate connectivity when a non-stationarity mean is present in the data. There will be little effect when changing this, but if it was reduced it will eventually be identical to simulation 2. The non-stationarity introduced by the adding the HRF function to the data is shown in Figure 7.

The HRF lasting 20 seconds with an assumed TR of 2 means that there are 40 seconds between trials. This is quite long for an event related fMRI experiment. If this was reduced, it would just entail that the HRF stacks due to overlap. This would have no effect on what the simulation is testing which is just how well the different methods can act upon non-stationarities. The TR was considered to be 2 seconds because many studies still use this TR despite lower TRs being available. However, this has no implications on the results.

### Assumptions regarding shift in “brain state”

The duration of a state in the quick condition are on the approximate time scale found in (4), although slightly quicker (where the average state transition was 7 seconds). The longer transitions are based on the time scale explored in (5). Note however that these two different time scales for states originate from different dynamic functional connectivity methods. Detecting brain states has, to our knowledge, not been justified without using dynamic functional connectivity which makes the reasoning to justify this parameter somewhat circular. Simulation 4 was added primarily to evaluate how the different methods can capture dynamics that many researchers hypothesize dynamic functional connectivity to be, not necessarily how the data is.

### Caveats

The current simulations do not take into account any frequency information of fMRI brain connectivity which is known to play a role in resting state fMRI. This means that any methods that utilize this aspect of the signal (e.g. phase) cannot be evaluated at the moment. The Gaussian assumption of the distributions can be improved upon by adding more fMRI specific noise. Moreover, negative correlations were not considered in the simulations.

The autocorrelation of *r*_*t*_ in simulations 2 and 3 do not necessarily entail an autocorrelation of *X*. While this could in principle be added to the simulation model, it would mean there is a greater correlation between the time series in *X* that is not known by the parameter *r*_*t*_. This caveat does not however impact the connectivity between the two signals.

## References

1. Hutchison RM, Hashemi N, Gati JS, Menon RS, Everling S. Electrophysiological signatures of spontaneous BOLD fluctuations in macaque prefrontal cortex. NeuroImage. 2015;113:257–67.

2. Ma S, Calhoun VD, Phlypo R, Adali T. Dynamic changes of spatial functional network connectivity in healthy individuals and schizophrenia patients using independent vector analysis. NeuroImage. 2014 Apr;90:196–206.

3. Kaiser RH, Whitfield-Gabrieli S, Dillon DG, Goer F, Beltzer M, Minkel J, et al. Dynamic Resting-State Functional Connectivity in Major Depression. Neuropsychopharmacology. 2015;1–9.

4. Betzel RF, Satterthwaite TD, Gold JI, Bassett DS. Positive affect, surprise, and fatigue are correlates of network flexibility. Scientific Reports. 2017;7(1):520.

5. Kucyi A, Hove MJ, Esterman M, Hutchison RM, Valera EM. Dynamic Brain Network Correlates of Spontaneous Fluctuations in Attention. Cerebral cortex (New York, NY: 1991). 2016;bhw029.

6. Barttfeld P, Uhrig L, Sitt JD, Sigman M, Jarraya B, Dehaene S. Correction for Barttfeld et al., Signature of consciousness in the dynamics of resting-state brain activity. Proceedings of the National Academy of Sciences. 2015;112(37):E5219–20.

7. Thompson WH, Fransson P. On Stabilizing the Variance of Dynamic Functional Brain Connectivity Time Series. Brain Connectivity. 2016 Dec;6(10):735–46.

8. Thompson WH, Brantefors P, Fransson P. From static to temporal network theory – applications to functional brain connectivity. Network Neruoscience. 2017;1(2):1–37.

9. Betzel RF, Fukushima M, He Y, Zuo XN, Sporns O. Dynamic fluctuations coincide with periods of high and low modularity in resting-state functional brain networks. NeuroImage. 2016;127(February 2016):287–97.

10. Thompson WH, Fransson P. The mean–variance relationship reveals two possible strategies for dynamic brain connectivity analysis in fMRI. Frontiers in Human Neuroscience. 2015;9(398):1–7.

11. Thompson WH, Fransson P. Bursty properties revealed in large-scale brain networks with a point-based method for dynamic functional connectivity. Scientific Reports. 2016 Dec;6(November):39156.

12. Laumann TO, Snyder AZ, Mitra A, Gordon EM, Gratton C, Adeyemo B, et al. On the Stability of BOLD fMRI Correlations. Cerebral Cortex. 2016;1–14.

13. Zalesky A, Breakspear M. Towards a statistical test for functional connectivity dynamics. NeuroImage. 2015;114:466–70.

14. Hindriks R, Adhikari MH, Murayama Y, Ganzetti M, Mantini D, Logothetis NK, et al. Can sliding-window correlations reveal dynamic functional connectivity in resting-state fMRI? NeuroImage. 2016;127:242–56.

15. Allen EA, Damaraju E, Plis SM, Erhardt EB, Eichele T, Calhoun VD. Tracking whole-brain connectivity dynamics in the resting state. Cerebral cortex (New York, NY: 1991). 2014 Mar;24(3):663–76.

16. Shine JM, Koyejo O, Bell PT, Gorgolewski KJ, Gilat M, Poldrack RA. Estimation of dynamic functional connectivity using Multiplication of Temporal Derivatives. NeuroImage. 2015;122:399–407.

17. Liu X, Duyn JH. Time-varying functional network information extracted from brief instances of spontaneous brain activity. Proceedings of the National Academy of Sciences of the United States of America. 2013 Mar;110(11):4392–7.

18. Leonardi N, Richiardi J, Gschwind M, Simioni S, Annoni J-M, Schluep M, et al. Principal components of functional connectivity: a new approach to study dynamic brain connectivity during rest. NeuroImage. 2013 Dec;83:937–50.

19. Tagliazucchi E, Balenzuela P, Fraiman D, Chialvo DR. Criticality in largescale brain FMRI dynamics unveiled by a novel point process analysis. Frontiers in physiology. 2012 Jan;3(February):15.

20. Tagliazucchi E, Siniatchkin M, Laufs H, Chialvo DR. The Voxel-Wise Functional Connectome Can Be Efficiently Derived from Co-activations in a Sparse Spatio-Temporal Point-Process. Frontiers in Neuroscience. 2016 Aug;10(AUG):1–13.

21. Kang J, Wang L, Yan C, Wang J, Liang X, He Y. Characterizing dynamic functional connectivity in the resting brain using variable parameter regression and Kalman filtering approaches. NeuroImage. 2011;56(3):1222–34.

22. Molenaar PCM, Beltz AM, Gates KM, Wilson SJ. State space modeling of time-varying contemporaneous and lagged relations in connectivity maps. NeuroImage. 2016;125:791–802.

23. Liao W, Wu G-R, Xu Q, Ji G-J, Zhang Z, Zang Y-F, et al. DynamicBC: A MATLAB Toolbox for Dynamic Brain Connectome Analysis. Brain Connectivity. 2014 Dec;4(10):780–90.

24. Smith SM, Miller KL, Moeller S, Xu J, Auerbach EJ, Woolrich MW, et al. Temporally-independent functional modes of spontaneous brain activity. Proceedings of the National Academy of Sciences of the United States of America. 2012 Feb;109(8):3131–6.

25. Kiviniemi V, Vire T, Remes J, Elseoud AA, Starck T, Tervonen O, et al. A sliding time-window ICA reveals spatial variability of the default mode network in time. Brain connectivity. 2011 Jan;1(4):339–47.

26. Lindquist MA, Xu Y, Nebel MB, Caffo BS. Evaluating dynamic bivariate correlations in resting-state fMRI: A comparison study and a new approach. NeuroImage. 2014;101:531–46.

27. Senden M, Reuter N, Heuvel MP van den, Goebel R, Deco G. Cortical rich club regions can organize state-dependent functional network formation by engaging in oscillatory behavior. NeuroImage. 2017;146(October 2016):561–74.

28. Ou J, Xie L, Jin C, Li X, Zhu D, Jiang R, et al. Characterizing and Differentiating Brain State Dynamics via Hidden Markov Models. Brain Topography. 2015 Sep;28(5):666–79.

29. Ryali S, Supekar K, Chen T, Kochalka J, Cai W, Nicholas J, et al. Temporal Dynamics and Developmental Maturation of Salience, Default and Central-Executive Network Interactions Revealed by Variational Bayes Hidden Markov Modeling. PLOS Computational Biology. 2016;12(12):e1005138.

30. Salvatier J, Wiecki TV, Fonnesbeck C. Probabilistic programming in Python using PyMC3. PeerJ Computer Science. 2016 Apr;2:e55.

31. Van Der Walt S, Colbert SC, Varoquaux G. The NumPy array: A structure for efficient numerical computation. Computing in Science and Engineering. 2011;13(2):22–30.

32. Oliphant TE. SciPy: Open source scientific tools for Python. Computing in Science and Engineering. 2007;9:10–20.

33. Hunter JD. Matplotlib: A 2D graphics environment. Computing in Science and Engineering. 2007;9(3):99–104.

34. Waskom M, Botvinnik O, Drewokane, Hobson P, David, Halchenko Y, et al. seaborn: v0.7.1 (June 2016). doiorg.

35. Leonardi N, Van De Ville D. On spurious and real fluctuations of dynamic functional connectivity during rest. NeuroImage. 2015 Sep;104(1):430–6.

36. Richter CG, Thompson WH, Bosman CA, Fries P. A jackknife approach to quantifying single-trial correlation between covariance-based metrics undefined on a single-trial basis. NeuroImage. 2015;114(January 2016):57–70.

37. Hoffman M, Gelman A. The No-U-Turn Sampler: Adaptively Setting Path Lengths in Hamiltonian Monte Carlo. Journal of Machine Learning Research. 2014;15:30.

38. Friston KJ, Holmes aP, Worsley KJ, Poline J-P, Frith CD, Frackowiak RSJ. Statistical parametric maps in functional imaging: A general linear approach. Human Brain Mapping. 1995;2(4):189–210.

39. Shakil S, Lee CH, Keilholz SD. Evaluation of sliding window correlation performance for characterizing dynamic functional connectivity and brain states. NeuroImage. 2016;133:111–28.

## Supplementary References

1. Gordon EM, Laumann TO, Gilmore AW, Petersen SE, Nelson SM, Dosenbach NUF, et al. Precision Functional Mapping of Individual Human NeuroResource Precision Functional Mapping of Individual Human Brains. Neuron. 2017;1–17.

2. Shen X, Tokoglu F, Papademetris X, Constable R. Groupwise whole-brain parcellation from resting-state fMRI data for network node identification. NeuroImage. 2013 Nov;82(October 2009):403–15.

3. Thompson WH, Fransson P. On Stabilizing the Variance of Dynamic Functional Brain Connectivity Time Series. Brain Connectivity. 2016 Dec;6(10):735–46.

4. Thompson WH, Fransson P. Bursty properties revealed in large-scale brain networks with a point-based method for dynamic functional connectivity. Scientific Reports. 2016 Dec;6(November):39156.

5. Allen EA, Damaraju E, Plis SM, Erhardt EB, Eichele T, Calhoun VD. Tracking whole-brain connectivity dynamics in the resting state. Cerebral cortex (New York, NY: 1991). 2014 Mar;24(3):663–76.

